# *Trichoderma afroharzianum* T22 induces rhizobia and flavonoid-driven symbiosis to promote tolerance to alkaline stress in garden pea

**DOI:** 10.1101/2024.07.11.603139

**Authors:** Asha Thapa, Md Rokibul Hasan, Ahmad H. Kabir

## Abstract

Soil alkalinity is a limiting factor for crops, yet the role of beneficial fungi in mitigating this abiotic stress in garden pea is understudied. In this study, *Trichoderma afroharzianum* T22 colonized the roots of garden pea cultivars exposed to soil alkalinity in a host-specific manner. In alkaline-exposed Sugar Snap, T22 improved growth parameters, consistent with increased tissue mineral content, particularly Fe and Mn, as well as enhanced rhizosphere siderophore levels. The split-root assay demonstrated that the beneficial effects of T22 on alkaline stress mitigation are the result of a whole-plant association rather than localized root-specific effects. RNA-seq analysis showed 575 and 818 differentially expressed genes upregulated and downregulated in the roots inoculated with T22 under alkaline conditions. The upregulated genes were mostly involved in the flavonoid biosynthetic pathway (monooxygenase activity, ammonia-lyase activity, 4-coumarate-CoA ligase), along with genes related to mineral transport and redox homeostasis. Further, a flavonoid precursor restored plant health even in the absence of T22, confirming the role of microbial symbiosis in mitigating alkaline stress. Interestingly, T22 restored the abundance of rhizobia, particularly *Rhizobium leguminosarum* and *Rhizobium indicum*, along with the induction of *NifA*, *NifD*, and *NifH* in nodules, suggesting a connection between T22 and rhizobia under soil alkalinity. Further, the elevated rhizosphere siderophore, root flavonoid, expression of *PsCoA* (4-coumarate-CoA ligase) as well as the relative abundance of *TaAOX1* and *R. leguminosarum* diminished when T22 was substituted with exogenous Fe. This suggests that exogenous Fe eliminates the need for microbiome-driven mineral mobilization, while T22-mediated alkaline stress mitigation depends on flavonoid-driven symbiosis and *R. leguminosarum* abundance. It was further supported by the positive interaction of T22 on *R. leguminosarum* growth in alkaline media. Thus, the beneficial effect of T22 on rhizobia likely stems from their interactions, not solely from the improved mineral status, particularly Fe, in plants. This study provides the first mechanistic insights into T22 interactions with host and rhizobia, advancing microbiome strategies to alleviate soil alkalinity in peas and other legumes.

## 1 Introduction

Soil alkalinity can result from elevated levels of bicarbonate and salts, increased cation exchange capacity, and reduced organic matter content (Zhao et al. 2020; Alcántara et al. 2000; Kroh and Pilon 2019). Soil alkalinity alters soil properties (Blaskó 2011) and can reduce micronutrient accumulation, leading to chlorosis, impaired photosynthesis, redox imbalance, and reduced crop yields (Kabir et al. 2022; Ferrarezi et al. 2022; Sun et al. 2016). Particularly, alkali-mediated iron (Fe) deficiency in legume crops can be severe, primarily because of the high levels of Fe required for nodule formation (Brear et al. 2013). Fe deficiency can influence both the legume host and the rhizobia individually, or it can directly affect their interactions (Brear et al. 2013; Dixon and Kahn 2004). It has been observed that *Rhizobium leguminosarum* is a prevalent type of rhizobia capable of forming nitrogen-fixing nodules on pea roots (Boeglin et al. 2022).

Different plants employ various strategies for Fe uptake, which are often disrupted under alkaline stress. Dicot plants utilize what is known as Strategy I for Fe acquisition (Waters et al. 2018; Curie and Briat 2003). Under alkaline conditions, Strategy I plants typically possess ferric chelate reductase activity, which reduces Fe(III) to the more soluble Fe(II) form (Kabir et al. 2012). Plants also acidify the rhizosphere through H secretion, a process that helps counteract high soil pH and enhances the solubility of not only Fe but also other micronutrients (Molnár et al. 2023; Kabir et al. 2012). Additionally, phenolic compounds secreted through root exudates chelate micronutrients, aiding their mobilization under alkaline stress (Curie and Mari 2017; Rodríguez-Celma and Schmidt 2013). Once solubilized, Fe is transported into the root via metal transporters such as *IRT1* (Fe-regulated transporter), which are common in many dicot species (Kabir et al. 2012; Vert et al. 2001). However, the efficiency of these responses under alkaline stress is often genotype-dependent, making many high-yielding cultivars particularly susceptible to alkaline stress.

Plants can accommodate a vast array of microorganisms in and around their organs which may offer a sustainable solution for fortifying crops with nutrient supplies under alkaline stress (Kabir and Bennetzen 2024; Jin et al. 2014). Among the beneficial fungi, arbuscular mycorrhizal fungi (AMF) appear to be almost exclusively studied for their helpful role in plants. However, *Trichoderma* spp. is one of the main groups of rhizosphere mycorrhizal fungi, other than AMF, that can promote plant growth and development through a variety of means (Harman et al. 2004; Qi and Zhao 2013). Of them, *T. afroharzianum* T22 has already demonstrated shreds of evidence for plant growth promotion and abiotic stress tolerance (Staropoli et al. 2024; Bjørk et al. 2023; Kabir et al. 2022). Also, several *Trichoderma* species are known to play a role as biocontrol agents by exhibiting strong antagonistic activity against plant pathogens (Singh et al. 2024; Ren et al. 2022). However, there are undoubtedly still a lot of unanswered questions regarding the mechanisms of connection with this fungus with host plants in response to biotic and abiotic stresses. In a previous study, we reported that T22 triggered Strategy-I responses in alkali-mediated Fe-deficient soybean, leading to the upregulation of *GmFRO2* and *GmIRT1* (Kabir et al. 2022). Recent research has shown that T22 significantly upregulated multiple genes related to Fe uptake, auxin synthesis, nicotianamine synthase 3, and phytosiderophore transporter 1, aiding sorghum plants in coping with indirect Fe deficiency (Kabir and Bennetzen 2024). Studies suggest that fungi-plant symbiosis is involved in local and/or systemic signaling, predominantly depending on host genetic factors (Giovannetti et al. 2024; MacLean et al. 2017). Symbiosis requires continual signaling between the symbionts and the activation of new transcriptional programs in the plant host, especially in the roots (Bucher et al. 2014). This process triggers changes essential for the plant cell to host the fungal endosymbiont and maintain symbiosis. Of many molecules involved in symbiosis, phytohormones play a key role in the symbiosis between most land plants and fungi (Liao et al. 2018; Lopez-Obando et al. 2015). Other than plant hormones, flavonoids released by plants, have been found to promote microbial colonization and serve as pre-symbiotic signaling molecules in microbial-plant symbiosis (Lidoy et al. 2023; Schliemann et al. 2008). Coumarins, secondary metabolites synthesized via the phenylpropanoid pathway in plants, act as signaling molecules regulating interactions between symbiotic microbes and plants (Harbort et al. 2020; Stringlis et al. 2009). However, this information is still limited to very few crops.

Garden pea (*Pisum sativum* ssp. *hortense*) is an important pulse crop with significant nutritional value, genetic diversity, and available genetic tools, making it a valuable research species (Pandey et al. 2022; Hacisalihoglu et al. 2021; Chaudhari et al. 2020). Addressing alkaline stress in crops often involves the use of costly and environmentally unsafe soil amendments and fertilizers. Evidence suggests that microbiomes can positively influence mineral status in plants (Kabir and Bennetzen 2024; Kabir et al. 2022; Bhardwaj et al. 2014). However, the specific role and mechanistic basis of T22 in pea plants, particularly its impact on nodule formation under alkaline stress, remains unclear. Therefore, we investigated whether T22 colonizes garden peas in a host-specific manner and how it induces transcriptional changes that promote symbiotic signaling and alkaline stress tolerance. Additionally, we sought to determine if the restoration of rhizobia in root nodules is due to the direct effect of T22 on rhizobia or its indirect effect through improving Fe availability in alkali-exposed peas.

## 2 Materials and methods

### 2.1 Plant cultivation and growth conditions

Seeds from six garden pea cultivars (Lillian, Little Marvel, Green Arrow, Sugar Sprint, Sugar Bon, and Sugar Snap) were used in this study. Based on initial screening, Sugar Snap was selected for further experiments. Seeds were surface sterilized by soaking in a 4% sodium hypochlorite solution for 5 min, followed by three washes with sterile water. They were then placed in a tray and incubated at 25°C for 2 d. After germination, uniform and healthy seedlings were transplanted into pots (containing 500 g of a soil mix) comprising soil collected from a pea field and commercial potting mix in a 1:2 ratio in four different treatments of lime (7.5 g of NaHCO_3_ and 4.5 g of CaCO_3_) and *Trichoderma afroharzianum* T22 (1 ×10 cfu/gram) as follows: (i) control (soil without lime pH ∼6.5), (ii) alkaline soil (soil mixed with lime pH ∼7.8), (iii) alkaline soil + T22 (soil mixed with lime pH ∼7.8 and T22 inoculum and (iv) T22+ (soil without lime pH ∼6.5 and T22 inoculum. The T22 inoculum was cultured on potato dextrose agar (PDA) medium to verify the viability of the fungal spores before inoculation (Supplementary Fig. S1). During the cultivation period, 15 mM NaHCO_3_ was added weekly to maintain the pH of the alkaline soil at ∼7.8. This cultivation process continued for 4 weeks prior to data measurement.

In a targeted experiment, cinnamic acid, a precursor in the flavonoid biosynthetic pathway (Li et al. 2020), was supplemented into the soil (0.5 g/500 g soil) to assess its role in inducing alkaline stress tolerance without exogenous T22. Additionally, two new treatment groups were included: FeEDDHA+ (soil mixed with lime, pH ∼7.8, and FeEDDHA at 0.5 g/500 g soil) and control (soil without lime, pH ∼6.5, and FeEDDHA at 0.5 g/500 g soil). These targeted studies were conducted for 5 weeks before data analysis. All experiments were performed in a randomized complete block design with 9 biological replications per treatment in a growth chamber with a light regimen of 10 h of light and 14 h of darkness (250 μmol m^2^ s^-1^) at approximately 25°C. The physicochemical characteristics of the soil are detailed in Supplementary Table S1.

### 2.2 Measurement of morphological features and chlorophyll fluorescence kinetics

The shoot height and root length of each plant were measured using a measuring tape. The harvested shoots and roots were then dried for 3 d at 70°C in an electric oven before recording their dry weights. For nodule assessment, the roots were gently washed to remove soil particles, and nodules were carefully harvested, counted with a magnifying lens, and stored at -80°C for further analysis. Additionally, chlorophyll content and fluorescence kinetics (OJIP), including Fv/Fm (maximal photochemical efficiency of PSII) and Pi_ABS (photosynthetic performance index), were measured on the uppermost fully developed leaves at three different spots. Chlorophyll content was assessed using a handheld SPAD meter (AMTAST, United States). The chlorophyll score refers to the relative measure of chlorophyll content in leaf tissue, obtained using a chlorophyll meter to assess the photosynthetic capacity and health in plants (Uddling et al. 2007). In addition, fluorescence kinetics were measured with a FluorPen FP 110 (Photon Systems Instruments, Czech Republic). For OJIP analysis, leaves were dark-adapted for 1 h before data collection.

### 2.3 Determination of H_2_O_2_ in roots

For H_2_O_2_ determination, root samples were mixed in a solution of 0.1% trichloroacetic acid, as previously described (Alexieva et al. 2001). Briefly, the samples were finely ground into a powder using a mortar and pestle. The extracted fluid underwent centrifugation for 15 min at 10,000 rpm to separate cellular residues. The resulting upper aqueous phase was supplemented with potassium iodide (1 M) and phosphate buffer (10 mM, pH adjusted to 7.0), and then left in darkness for 60 min. Finally, the optical density of the solution was measured at 390 nm using a spectrophotometer (ThermoFisher, Waltham, USA), and the concentration of H_2_O_2_ was calculated based on the standard curve.

### 2.4 Molecular detection of T22 colonization in roots

To account for potential variation in T22 colonization across different root sections of garden pea, we performed root partitioning to identify the segment with the highest colonization. The roots were divided into four segments: the mature zone (1–3 cm), the elongation zone (3–6 cm), the meristematic zone (6 cm and beyond), and the root tips (Supplementary Fig. S4). The establishment and colonization of T22 in the roots of Sugar Snap cultivated under alkaline stress were evaluated by measuring the relative abundance of a T22 marker gene (*TaAOX1*: L-amino acid oxidase gene) using DNA-based qPCR (Kabir and Bennetzen 2024). Briefly, root samples were washed twice in fresh sterile phosphate-buffered saline, followed by a brief vortex and two rinses with sterile water. The DNA was extracted on approximately 0.2 g of root samples using CTAB (cetyltrimethylammonium bromide) method (Clarke 2009). Subsequently, DNA was quantified using a NanoDrop ND-1000 Spectrophotometer (Wilmington, USA) and diluted to equal quantities for each DNA sample. The qPCR analysis was conducted using a CFX96 Touch Real-Time PCR Detection System (Bio-Rad, USA), with specific primers (forward: 5′-GTC GGT AGC TGA AAG GGG AT-3′; reverse: 5′-AT TAG AGG CCG GAA ACA CC-3′) designed to amplify a fragment of *TaAOX1* of T22 (Kabir and Bennetzen 2024). Additionally, the relative abundance of *Rhizobium leguminosarum* (RHIZ) was detected in the nodules using specific primer pairs (Boonen et al. 2010). The iTaq™ Universal SYBR® Green Supermix (Bio-Rad, USA) was used in all reactions along with gene-specific primers (Supplementary Table S2) for *PsGAPDH* that served as the internal control for relative quantification using 2-^ΔΔ^CT method (Livak and Schmittgen 2001).

### 2.5 Determinations of siderophore levels in the rhizosphere

Siderophore content in the rhizosphere soil was determined using a chrome azurol S (CAS) assay (Himpsl and Mobley 2019). Briefly, rhizosphere soil samples were homogenized with 80% methanol and centrifuged for 15 min at 10,000 rpm. The supernatant (500 μl) was mixed with 500 μl of CAS solution and incubated at room temperature for 5 min. Absorbance was measured at 630 nm, using 1 ml of CAS reagent as a reference. Siderophore levels per gram of soil sample were calculated with the formula: % siderophore unit = [(Ar - As) / Ar] x 100, where Ar is the absorbance of the reference (CAS reagent only) at 630 nm and As is the absorbance of the sample (CAS reagent + soil sample) at 630 nm.

### 2.6 Nutrient analysis in root and leaf

Nutrient analysis was performed on both roots and young leaves. The roots were cut, washed with running tap water, and then immersed in 0.1 mM CaSO_4_ for 10 min, followed by a rinse with deionized water. Young leaves were separately washed with deionized water. All samples were placed in individual envelopes and dried in an electric oven at 75°C for 3d. Elemental concentrations were measured using inductively coupled plasma mass spectrometry (ICP-MS) at the Agricultural and Environment Services Laboratories at the University of Georgia.

### 2.7 Split-root assay

The plants were cultivated for 2 weeks in sterile vermiculite before conducting the split root assay in soil pots. The split-root plants were cultivated for an additional four weeks before data collection. Briefly, split root was done using a single pot with a partitioned split-root assembly (compartment) with some modifications as previously described (Thilakarathna and Cope 2021). The taproot of each seedling was cut in the middle, and the lateral root system was evenly divided between two compartments within a single pot, filled with natural field soil and commercial potting mix (1:2). The compartments were separated by a solid plastic barrier. Each split-root system was subjected to the same growth conditions, including the presence or absence of soil alkalinity. In one compartment of each split-root system, T22 was inoculated, while the other compartment remained uninoculated, based on the treatment.

### 2.8 RNA-sequencing and bioinformatics analysis

RNA-seq analysis was conducted on roots collected from the meristematic zone. Before RNA extraction, roots were thoroughly cleaned by rinsing twice with sterile water and vortexing for 10 sec in sterile phosphate-buffered saline to remove surface contaminants. The cleaned roots were then ground into a powder using liquid nitrogen and a pre-cooled pestle and mortar. RNA extraction was performed using the SV Total RNA Isolation System (Promega Corporation, USA). One microgram of RNA per sample, with a RIN value higher than 8, was used for RNA-seq library preparation. Libraries were prepared with the KAPA RNA HyperPrep Kit with Poly-A Selection (Kapa Biosystems, USA), followed by amplification using the KAPA HiFi HotStart Library Amplification Kit (Kapa Biosystems, USA). Sequencing was carried out on an Illumina NovaSeq 6000 instrument with 0.2 Shared S4 150 bp paired-end sequencing at the RTSF Genomics Core at Michigan State University. A total of 92.5% of reads passed filters with a quality score of ≥ Q30, generating 917.23 Gbp of data.

Bioinformatics analysis of raw FastQ files was carried out using Partek Flow genomic analysis software (Partek, St. Louis, MO, USA). High-quality clean reads were obtained by removing adaptor sequences and low-quality reads using the Trimmomatic tool (Bolger et al. 2014). The clean reads were aligned to the pea reference genome using HISAT2 (Kim et al. 2015). Read counts mapped to each gene were quantified with HTSeq (Anders et al. 2015). Differentially expressed genes (DEGs) were identified using DESeq2 (v1.16.1) with the following criteria: an adjusted p-value (FDR) < 0.05 and a log fold change ≥ 2. Genes with low expression (<10 normalized counts across all samples) were excluded. This analysis was based on FPKM (fragments per kilobase of exon per million mapped fragments). Venn diagrams were created using VennDetail (Hinder et al. 2017), and heatmaps were generated using pheatmap (Hu et al. 2021) in the R programming environment. Gene enrichment analyses of garden pea DEGs were performed using ShinyGO 0.76.3 (Ge et al. 2020), with gene ontology data sourced from EnsemblPlants resources (Yates et al. 2022). For co-expression network analysis, WGCNA (Weighted Gene Co-expression Network Analysis) was performed on normalized expression data to identify gene modules associated with KEGG pathways, using a pathway significance cutoff of FDR < 0.1.

### 2.9 Nanopore-based 16S sequencing in root nodules

The nodules were first washed with running tap water, then surface-sterilized in a 3% sodium hypochlorite solution for 3 min and rinsed three times with sterile distilled water. Total DNA was extracted using the CTAB method as described by Clarke (2009). PCR reactions were conducted with 5 ng of DNA from each sample and 16S primers 27F (AGAGTTTGATCMTGGCTCAG) and 1492R (CGGTTACCTTGTTACGACTT) to amplify the near full-length bacterial 16S rRNA gene. Thermal cycling conditions included an initial denaturation at 95°C for 2 min, followed by 30 cycles of denaturation at 95°C for 30 sec, annealing at 60°C for 30 sec, and extension at 72°C for 2 min. A final extension at 72°C for 10 min and a hold at 4°C were performed. Amplicons from each sample were ligated to pooled barcoded reads using the 16S Barcoding Kit 24 V14 (Oxford Nanopore Technologies, Oxford, UK) for library construction. Sequencing was conducted on a MinION portable sequencer (Oxford Nanopore Technologies) using a Flongle flow cell (R10.4.1). MinION™ sequencing data (FASTQ files) were processed using EPI2ME software from Oxford Nanopore Technologies to produce pass reads with an average quality score > 9. Adapter and barcode sequences were trimmed using the EPI2ME Fastq Barcoding workflow (Oxford Nanopore Technologies). Principal coordinate analysis, alpha diversity, and relative abundance of the bacterial community were calculated using the R packages phyloseq and vegan. Non-parametric Kruskal–Wallis tests were used to identify differences in taxa abundance at a 5% significance level (McMurdie and Holmes, 2013).

### 2.10 Real-time qPCR analysis of genes

Before RNA extraction from pea nodules, surface contaminants were thoroughly removed by washing with PBS buffer. Briefly, RNA extraction and cDNA conversion were performed using the SV Total RNA Isolation System (Promega Corporation, USA) and GoScript™ Reverse Transcriptase (Promega Corporation, USA) kits, respectively. Subsequently, the expressions of *NifA* (nitrogenase Fe protein regulatory protein), *NifD* (nitrogenase FeMo protein alpha subunit), and *NifH* (nitrogenase FeMo protein beta subunit) associated with the nitrogen fixation process in nodules as well as the expression of *PsCoA* (*Psat2g188920*) gene in roots were studied using gene-specific primers (Supplementary Table S2) in CFX96 Real-Time system (Bio-Rad, USA). The qPCR reaction was conducted with an initial denaturation at 95°C for 3 min, followed by 40 cycles of denaturation at 95°C for 5 sec, annealing at 55°C for 30 sec, and extension at 60°C for 5 min. Expression levels of target genes were analyzed using the dd^-ΔCt^ method (Livak and Schmittgen 2001), with normalization to *PsGAPDH*.

### 2.11 Determination of total flavonoid in roots

The total flavonoid content was measured in pea roots as previously described (Piyanete et al. 2009). Briefly, 0.5 mL of a 70% ethanol extract from the roots was mixed with 2 mL of distilled water and 0.15 mL of a 5% NaNO_2_ solution. After incubating for 6 min, 0.15 mL of a 10% AlCl_3_ solution was added, and the mixture was left to stand for another 6 min. Then, 2 mL of a 4% NaOH solution was added. The mixture was diluted to a final volume of 5 mL with methanol and thoroughly mixed. After a 15-min incubation period, the absorbance was measured at 510 nm using a spectrophotometer against a blank. The total flavonoid concentration was expressed in milligrams per gram of dry weight, calculated using a quercetin standard curve.

### 2.12 Microbial co-culture method

We studied the interactions between T22 and *R. leguminosarum* on nutrient agar under Fe-sufficient and Fe-deficient conditions, supplemented with 15 mM NaHCO_3_ to raise the pH to 7.8. The microbial species (5 µl inoculum) were cultured in three different combinations: T22 alone, *R. leguminosarum* alone, and T22 + *R. leguminosarum*. The growth of the microbial species was measured after 7d of culture using ImageJ software (National Institutes of Health, USA). The radius of microbial colonies was measured at three points and then averaged.

### 2.13 Data analysis

The statistical significance of each variable was assessed using analysis of variance (ANOVA), with the treatment considered as the independent variable. Subsequently, Duncan’s Multiple Range Test (DMRT) was performed using SPSS Statistics 20.0 (IBM). Graphical presentations were created with GraphPad Prism 7 and the R package ggplot2.

## 3 Results

### 3.1 Effect of T22 on root colonization and morphological features

The colonization efficiency of T22 in roots and its effects on the growth parameters of different garden pea cultivars exposed to alkaline stress was evaluated. The relative abundance of *TaAOX1* (a marker of T22 colonization) did not differ significantly in the roots of Lillian, Little Marvel, Green Arrow, and Sugar Sprint, whether inoculated with T22 or not, in the presence or absence of alkali-mediated soil alkalinity (Supplementary Fig. S2A). In Sugar Bon and Sugar Snap, alkaline stress caused no significant changes in *TaAOX1* abundance compared to controls.

However, T22 addition under soil alkalinity resulted in a significant increase in *TaAOX1* abundance in these two cultivars. Furthermore, T22 inoculation in control plants significantly increased the relative abundance of *TaAOX1* in Sugar Bon compared to plants inoculated with T22 under alkaline stress. Under control conditions, Sugar Snap inoculated with T22 showed a significant decrease in *TaAOX1* compared to the plants supplemented with T22 under alkaline stress (Supplementary Fig. S2A). We found that Sugar Snap showed a significant reduction in SPAD scores in leaves under alkaline stress. However, SPAD scores significantly increased in the leaves of alkali-exposed Sugar Snap inoculated with T22 relative to plants exposed to soil alkalinity. Control plants inoculated with T22 had a similar SPAD score to controls in Sugar Snap (Supplementary Fig. S2B). Sugar Snap showed a significant decline in root length and root dry weight under alkaline stress compared to controls. However, T22 supplementation in both alkali-exposed and control plants led to a significant improvement in root length and root dry weight compared to alkali-exposed plants without T22 (Supplementary Fig. S2C-S2D). Lastly, Sugar Snap exhibited significant declines in both shoot height and shoot dry weight under alkaline stress compared to controls. Alkali-exposed Sugar Snap inoculated with T22 showed a significant increase in these parameters to those of plants solely cultivated under Fe shortage. T22 inoculated in Sugar Snap under control conditions showed similar shoot height and shoot dry weight to those of controls (Supplementary Fig. S2E-S2F).

In this study, alkali-exposed Sugar Bon inoculated with T22 showed no significant increase in leaf SPAD score, root length, or shoot height compared to plants grown solely under alkaline stress (Supplementary Fig. S2B, S2C, and S2E). However, the root and shoot dry weights of alkali-exposed Sugar Bon inoculated with T22 significantly increased compared to plants under alkaline stress alone (Supplementary Fig. S2D-S2F). Among the other genotypes of garden peas, T22 inoculation showed no significant improvement in leaf SPAD score, root length, root dry weight, shoot height, and shoot dry weight in Lillian, Little Marvel, Green Arrow, and Sugar Sprint exposed to alkaline stress compared to plants grown solely under soil alkalinity (Supplementary Fig. S2B-S2F).

### 3.2 Effect of T22 on photosynthetic parameters and rhizosphere siderophore

In this study, alkaline stress caused substantial retardation in both aboveground shoot height and belowground root length growth in Sugar Snap (Fig. 1A-1D). However, plants inoculated with T22, regardless of soil pH, exhibited significant increases in both aboveground and belowground growth parameters compared to plants cultivated solely under alkaline stress (Fig. 1A-1D). Furthermore, leaf chlorophyll fluorescence indices showed significant variations in response to the absence or presence of alkaline stress and T22 (Fig. 1E-1F). The Fv/Fm significantly declined in the leaves due to alkaline stress compared to controls. The addition of T22 to alkali-exposed plants led to a significant increase in Fv/Fm relative to plants solely cultivated with alkaline stress. Plants inoculated with T22 under control conditions showed similar Fv/Fm values compared to untreated controls (Fig. 1E). Additionally, Pi_ABS significantly decreased in response to alkaline stress compared to controls. However, the addition of T22, regardless of soil pH, resulted in a significant increase in Pi_ABS in the leaves compared to alkali-exposed plants (Fig. 1F).

**Fig. 1.**
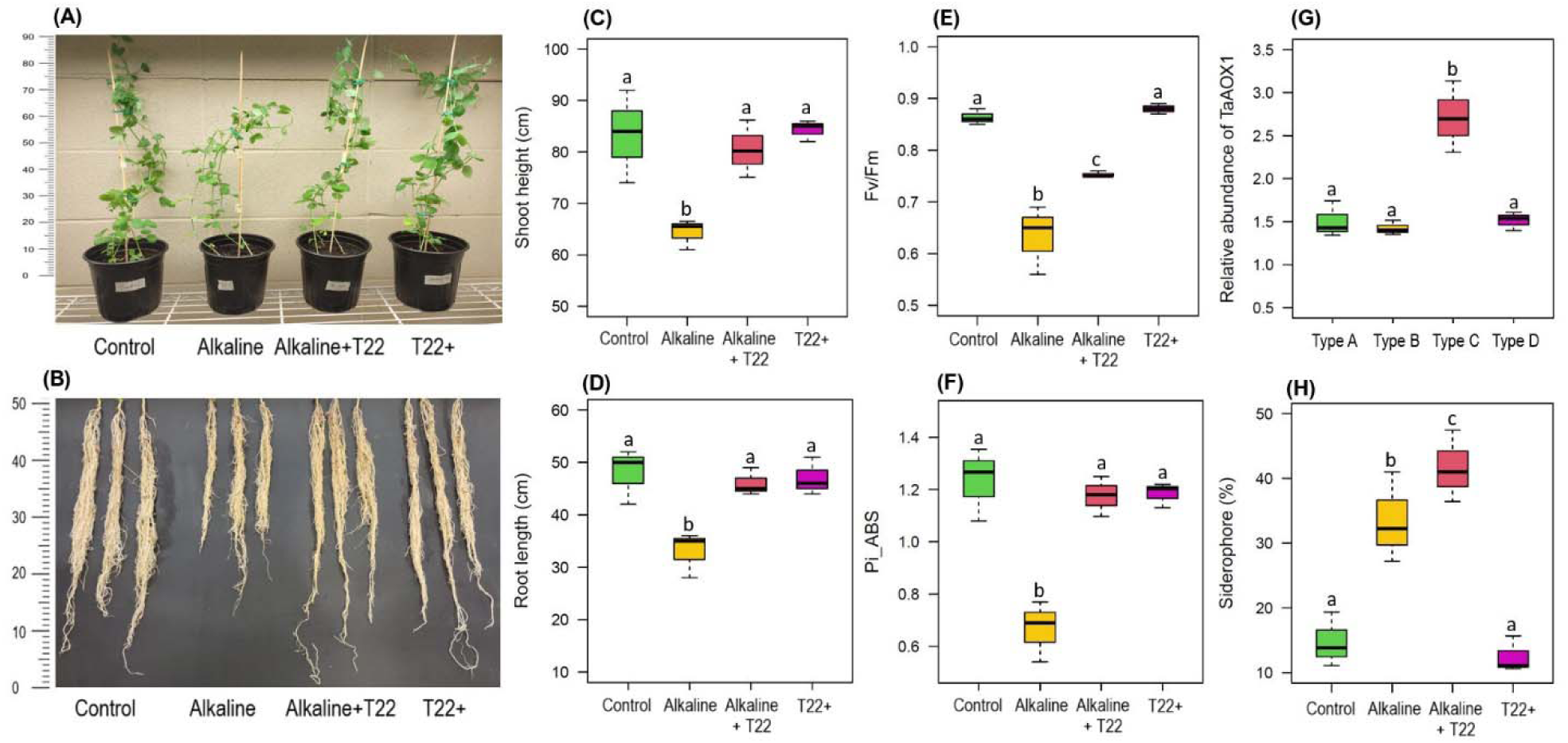
Changes in aboveground shoot phenotype (A), belowground root phenotype (B), shoot height (C), root length (D), leaf chlorophyll fluorescence indices: *Fv*/*Fm* -maximal quantum yield of PS II (E), Pi_ABS - photosynthetic performance index (F), the relative abundance of *TaAOX1* in different root partitions of Sugar Snap inoculated with T22 under alkaline stress (type A: mature zone, type B: elongation zone, type C: meristematic zone, type D: root tips) of Sugar Snap (G), and siderophore (%) in rhizosphere soil (H) in Sugar Snap cultivated in the absence or presence of alkaline stress inoculated with (T22+) or without T22. The data presented are means with standard deviations (*n* = 3 individual replicates). Statistically significant differences in the ANOVA test among different treatments (*p* < 0.05) are indicated by different letters above the bars, where applicable.

We also assessed the colonization efficiency of T22 in different root partitions of Sugar Snap plants cultivated under alkaline stress. The meristematic zone (type C) showed a significant increase in the relative abundance of *TaAOX1* compared to other root partitions (mature zone, elongation zone, root tips) of Sugar Snap (Fig. 1G). Further, alkaline stress led to a significant increase in siderophore levels in the rhizosphere compared to untreated controls (Fig. 1H). Interestingly, alkali-exposed plants inoculated with T22 exhibited a significant increase in siderophore levels in the rhizosphere soil compared to plants solely grown in soil alkalinity. The addition of T22 to untreated controls resulted in siderophore levels similar to controls, but significantly lower than those under alkali-exposed conditions, with or without T22 inoculation (Fig. 1H).

### 3.3 Changes in elemental content and hydrogen peroxide

During alkaline stress, Fe concentrations significantly decreased in both the roots and leaves of plants compared to Fe-sufficient controls (Table 1). Application of T22 to alkali-exposed plants significantly increased Fe levels in both roots and leaves compared to the plants grown solely under alkaline stress. Additionally, Zn and B levels showed a significant decrease in the roots in response to alkaline stress compared to untreated controls. Supplementation with T22 did not improve Zn and B levels in the roots of either alkali-exposed or untreated control plants. Further, Zn and B concentrations in the leaves remained unchanged across all treatment groups (Table 1). Mn levels significantly declined in both the roots and leaves of alkali-exposed plants compared to controls. However, alkali-exposed plants inoculated with T22 exhibited a significant increase in Mn levels compared to plants solely exposed to soil alkalinity. Also, untreated control plants inoculated with T22 showed equivalent Mn levels to those of controls (Table 1). Also, S content significantly decreased in both the roots and leaves of alkali-exposed plants compared to controls. However, no significant changes in S levels were observed when alkali-stressed plants were supplemented with T22. T22 inoculated in untreated control conditions showed similar S levels in roots, but the leaves showed a significant decrease in S levels compared to control plants (Table 1). Furthermore, no significant differences were found in Mg and P concentrations in both roots and leaves regardless of soil pH and T22 supplementation. Ca levels in the roots showed no significant changes across different treatment groups. However, Ca levels were significantly reduced in the leaves under alkali-exposed conditions relative to controls. Alkali-stressed plants supplemented with T22 showed a significant increase in Ca levels compared to plants solely cultivated under soil alkalinity. Untreated control plants inoculated with T22 had similar Ca levels in the leaves relative to controls (Table 1). Furthermore, hydrogen peroxide was significantly increased in the roots of plants exposed to alkaline conditions compared to untreated controls (Supplementary Fig. S3). However, alkali-exposed plants inoculated with T22 showed a significant decline in hydrogen peroxide compared to the plants solely cultivated under alkaline stress. Untreated plants inoculated with T22 showed similar hydrogen peroxide levels to those of controls (Supplementary Fig. S3).

**Table 1.**
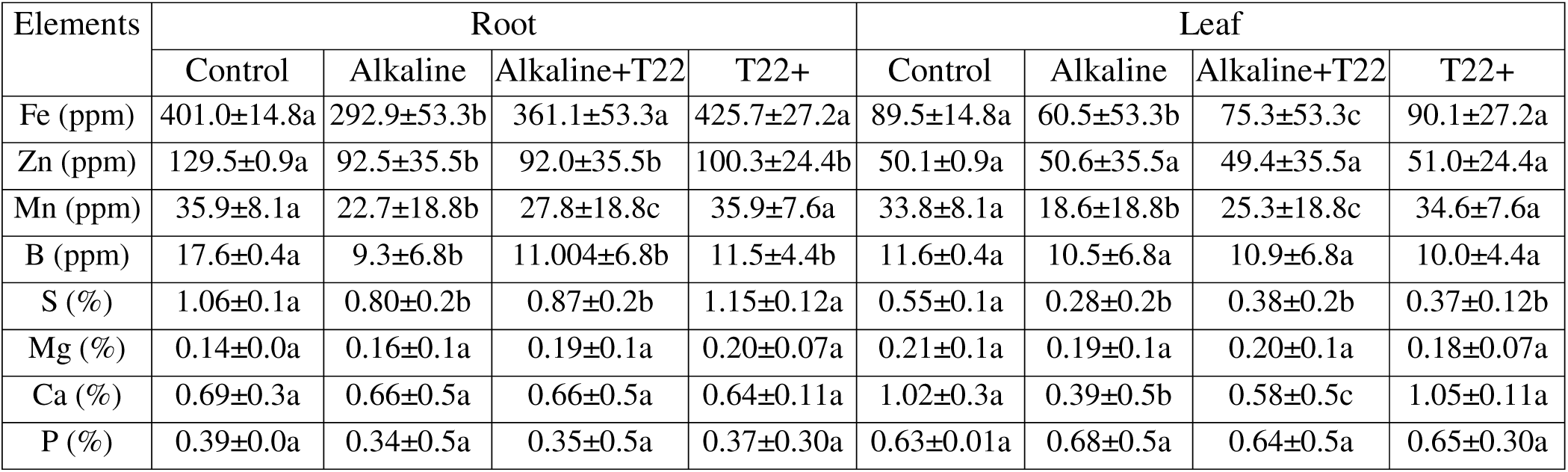
ICP-MS analysis of nutrients in the root and leaf of pea plants inoculated with or without *T. afroharzianum* (T22) in the absence or presence of alkaline stress. Data represents ± SD of three independent biological samples. Statistically significant differences in the ANOVA test among different treatments are indicated by different letters at the *p* < 0.05 level.

### 3.4 Determination of signaling cascade facilitating symbiotic association

We established a split-root system to investigate local versus systemic signaling in the symbiosis between T22 and garden pea (Sugar Snap) under different pH levels (Fig. 2A-2B). Plants cultivated without T22 in both compartments (-/-) showed significant reductions in shoot growth under soil alkalinity compared to controls (Fig. 2C). However, plants inoculated with T22 in both compartments (+/+) or in one compartment (-/+) exhibited significant improvements in shoot growth compared to alkali-imposed plants without T22 in any compartment (-/-). Interestingly, no phenotypic difference in shoot growth was observed between plants inoculated with T22 in both compartments (+/+) and those inoculated in just one compartment (-/+) (Fig. 2C). Similarly, plants cultivated without alkaline stress, whether inoculated with T22 in both compartments or just one, showed no significant differences in shoot phenotype compared to plants grown under control conditions (Fig. 2C). Alkali-stressed plants grown without T22 in both compartments (-/-) had markedly lower SPAD scores compared to those grown under control conditions (Fig. 2D). Plants inoculated with T22 in both compartments (+/+) had significantly higher SPAD scores in the leaves compared to alkali-exposed plants that were not inoculated with T22 in any compartment (-/-). However, plants inoculated with T22 in only one compartment (-/+) had relatively lower SPAD scores (P<0.05) compared to those inoculated with T22 in both compartments (+/+) (Fig. 2D). Lastly, plants cultivated under control conditions, whether inoculated with T22 in both (+/+) or just one (-/+) compartment, showed similar SPAD scores to plants solely cultivated under control conditions (Fig. 2D).

**Fig. 2.**
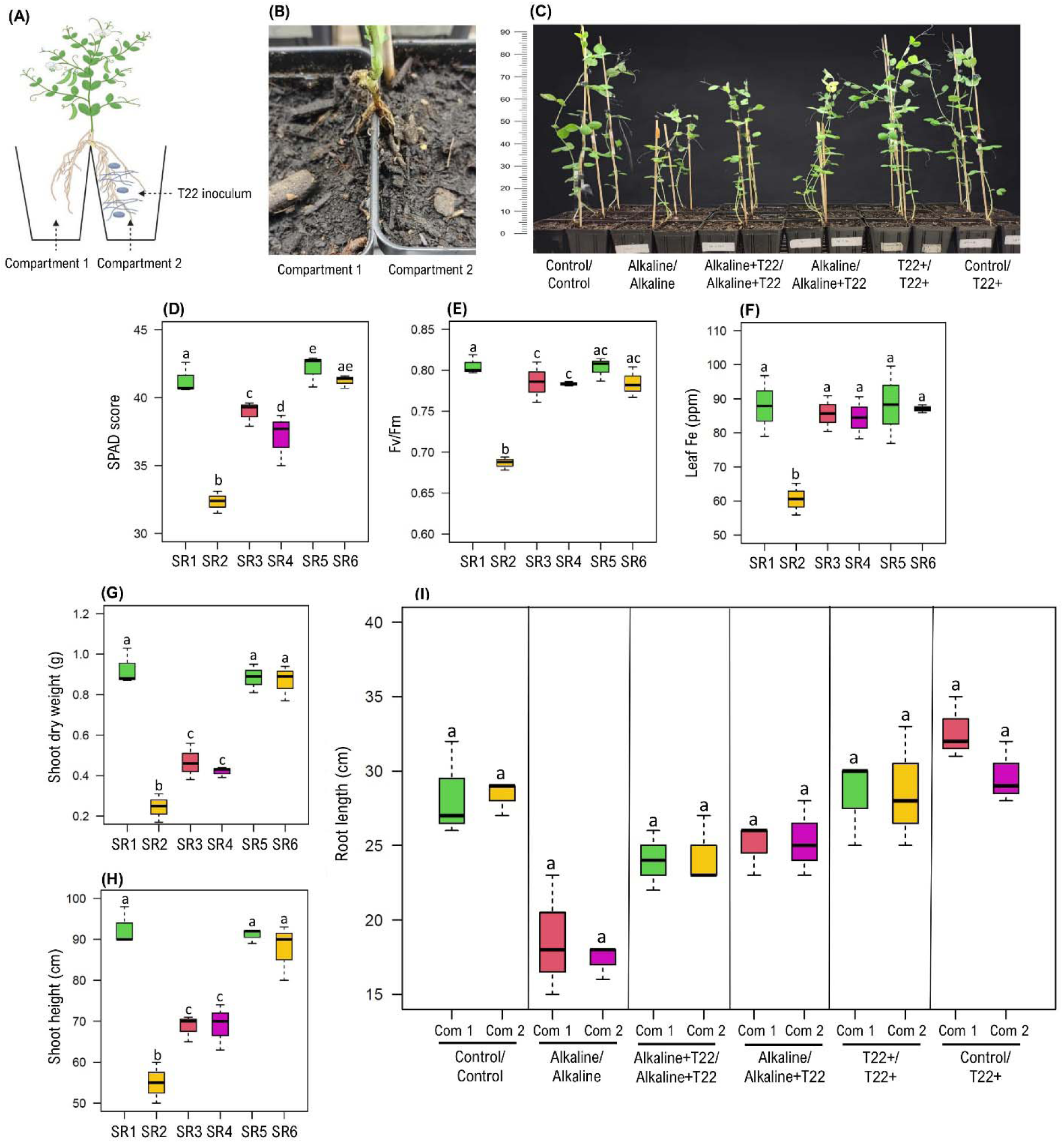
Schematic diagram of split-root (SR) system (A), establishment of split-root (B), plant phenotype (C), leaf SPAD score (D), leaf *Fv*/*Fm* - maximal quantum yield of PS II (E), leaf Fe (F), shoot dry weight (G), shoot height (H) and root length (I) of plants grown in each compartment in the absence (control) or presence (+) of alkaline stress and T22 inoculum. The data presented are means with standard deviations (*n* = 3 individual replicates). Statistically significant differences in the ANOVA test among different treatments (*p* < 0.05) are indicated by different letters above the bars, where applicable. SR1: control/control, SR2: alkaline/alkaline, SR3: alkaline+T22+/ alkaline+T22, SR4: alkaline/ alkaline+T22, SR5: T22+/T22+, SR6: Control/T22+.

Plants grown under alkaline stress without T22 in both compartments (-/-) had significantly lower Fv/Fm levels in their leaves compared to the plants that were cultivated in control conditions (Fig. 2E). Alkali-exposed plants inoculated with T22 in both (+/+) or any (-/+) compartment showed a significant increase in SPAD scores relative to alkali-stressed plants not inoculated with T22 in any compartment (Fig. 2E). In contrast, plants cultivated under control conditions and inoculated with T22 T22 in either both (+/+) or one (-/+) compartment exhibited similar Fv/Fm to those of plants not inoculated with T22 in control conditions (Fig. 2E). Further, Fe concentration in the leaves showed a significant decrease in plants grown under alkaline stress relative to plants cultivated without soil alkalinity (Fig. 2F). Interestingly, T22 inoculated in both (+/+) or any (-/+) of the compartment regardless of the soil pH showed a significant increase in leaf Fe levels compared to the alkali-exposed plants not inoculated with T22 (Fig. 2F). Additionally, plants grown under alkaline stress without T22 in either compartment (-/-) showed significant decreases in shoot dry weight and shoot height compared to controls (Fig. 2G-2H). However, the addition of T22 to either both (+/+) or a single (-/+) compartment under soil alkalinity significantly improved shoot dry weight and shoot height compared to alkali-exposed plants not inoculated with T22. Similarly, T22 inoculated in either both (+/+) or one (-/+) compartment without soil alkalinity showed similar shoot dry weight and shoot height to those of control plants (Fig. 2G-2H). Lastly, root length did not show significant changes regardless of the exogenous supply of T22 in different compartments in the presence or absence of alkaline stress (Fig. 2I).

### 3.5 Effect of T22 on transcriptome and gene enrichment under alkaline stress

We performed RNA-seq analysis using the meristematic segments of roots to explore the biological mechanisms by which T22 alleviates alkaline stress in Sugar Snap. Principal component analysis (PCA) revealed that the first principal component (PC1) accounted for 43.2% of the variance, while the second principal component (PC2) explained 15.9% of the variance (Fig. 3A), indicating substantial variability among the sample groups (Fig. 3A, Supplementary Fig. S5). The RNA-seq analysis demonstrated significant changes in the root transcriptome (fold-change threshold ≥2, p ≤ 0.05) in response to alkaline stress, with 364 genes upregulated and 478 genes downregulated (Fig. 3B and 3C). Compared to alkali-exposed plants, untreated plants inoculated with T22 showed significant upregulation of 575 genes and downregulation of 818 genes. Between alkaline+T22 and alkali-exposed conditions, only 3 overlapping genes were differentially upregulated, while 60 genes were differentially downregulated in the roots. In plants without soil alkalinity inoculated with T22, 417 genes were significantly upregulated, and 379 genes were downregulated. Of these, 54 genes were shared in the upregulated group and 165 genes in the downregulated group. Furthermore, 442 genes were significantly upregulated, and 367 genes were significantly downregulated in plants inoculated with T22 under control conditions compared to those under alkaline stress (Fig. 3B-3C).

**Fig. 3.**
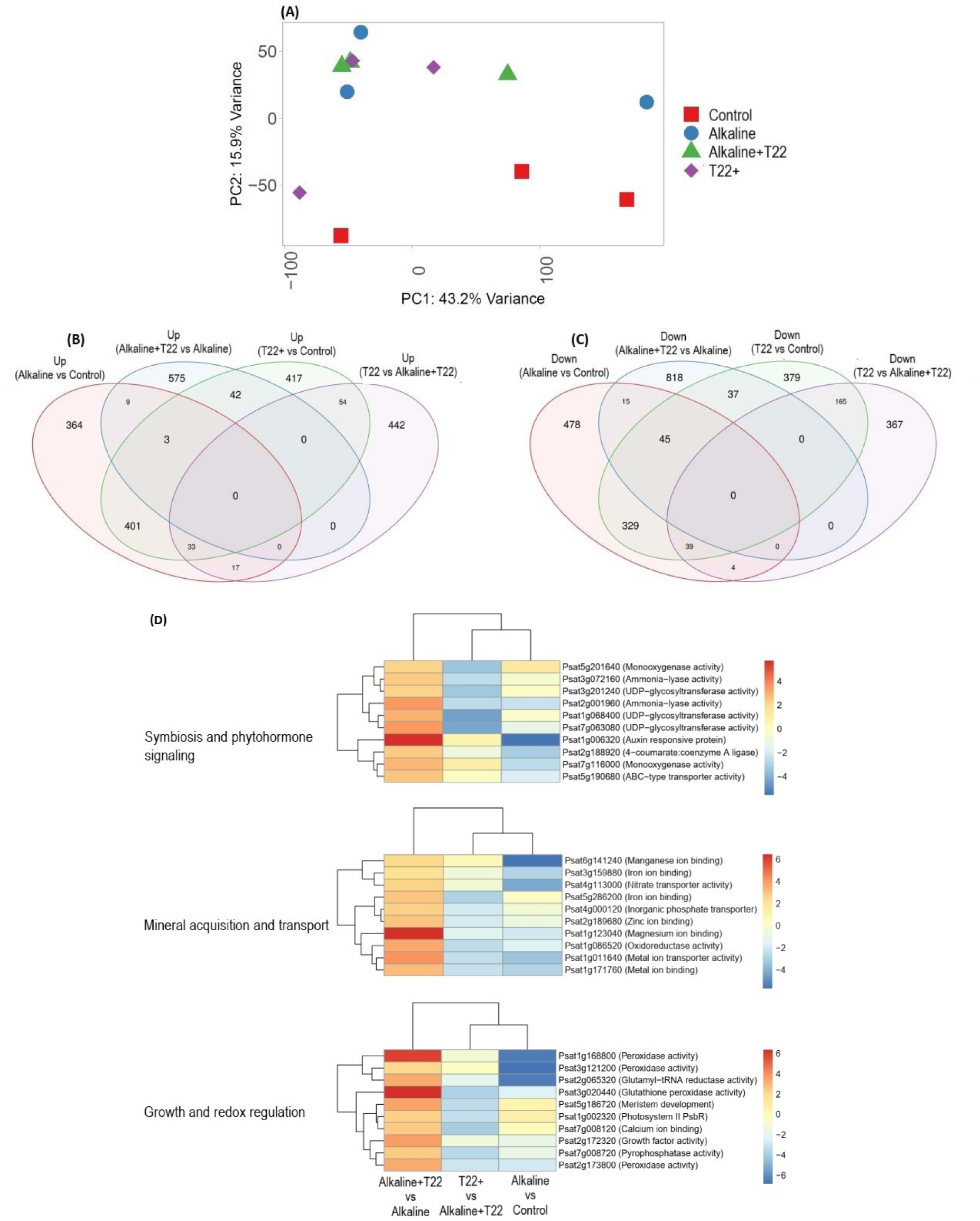
RNA-seq analysis: principal component analysis of gene variation (A), Venn diagram of differentially expressed (> Log2 fold, < P value 0.05) upregulated (B) and downregulated genes (C) and heatmap of top 10 upregulated genes (D) associated with different functions (symbiosis and phytohormone signaling, mineral acquisition and transport and growth and redox regulations) in the absence or presence of alkaline stress inoculated with (T22+) or without T22.

We further annotated the functions of DEGs using the Ensembl Genomes 58 database for the pea genome and summarized the top candidate genes due to the presence or absence of soil alkalinity and T22 inoculum (Fig. 3D, Supplementary Table S3). Among the genes associated with symbiosis and phytohormone signaling, the expression of *Psat5g201640* (monooxygenase activity), *Psat3g072160* (ammonia-lyase activity), *Psat3g201240* (UDP-glycosyltransferase activity), *Psat2g001960* (ammonia-lyase activity), *Psat1g068400* (UDP-glycosyltransferase activity), *Psat7g063080* (UDP-glycosyltransferase activity), *Psat1g006320* (auxin-responsive protein), *Psat2g188920* (4-coumarate A ligase), *Psat7g116000* (monooxygenase activity), and *Psat5g190680* (ABC-type transporter activity) showed significant upregulation in the roots of alkali-exposed plants inoculated with T22 compared to controls. However, some of these genes either remained unchanged or were significantly downregulated (*Psat5g201640*, *Psat3g072160*, *Psat3g201240*, *Psat1g068400*, *Psat7g063080*) under alkaline stress compared to controls (Fig. 3D, Supplementary Table S3).

In this study, several genes related to mineral acquisition and transport were significantly upregulated in alkali-stressed plants inoculated with T22 relative to plants solely grown under soil alkalinity which include: *Psat6g141240* (manganese ion binding), *Psat3g159880* (Fe ion binding), *Psat4g113000* (nitrate transporter activity), *Psat5g286200* (Fe ion binding), *Psat4g000120* (inorganic phosphate transporter), *Psat2g189680* (Zn ion binding), *Psat1g123040* (magnesium ion binding), *Psat1g086520* (oxidoreductase activity), *Psat1g011640* (metal ion transporter activity), *Psat1g171760* (metal ion binding). Most of these transcriptomes showed significant downregulation in the roots of plants exposed to alkaline stress relative to controls (Fig. 3D, Supplementary Table S3). We also found that the expression of *Psat1g168800* (peroxidase activity), *Psat3g121200* (peroxidase activity), *Psat2g065320* (glutamyl−tRNA reductase activity), *Psat3g020440* (glutathione peroxidase activity), *Psat5g186720* (meristem development), *Psat5g198360* (glutathione S-transferase), *Psat7g008120* (calcium ion binding), *Psat2g172320* (growth factor activity), *Psat7g008720* (pyrophosphatase activity), *Psat2g173800* (peroxidase activity) which are related to growth and redox regulation significantly induced in roots of Fe-deficient plants inoculated with T22 in contrast to Fe-deprived plants (Fig. 3D). While *Psat2g065320* (glutamyl-tRNA reductase activity), *Psat3g020440* (glutathione S-transferase), *Psat5g186720* (meristem development), *Psat1g002320* (growth factor activity), *Psat7g008120* (calcium ion binding), *Psat7g008720* (pyrophosphatase activity) and *Psat2g173800* (peroxidase activity) showed significant downregulation, the rest of the genes showed no significant changes in response to alkaline stress compared to controls (Fig. 3D). Furthermore, there were no significant changes in the expression levels of any of top genes discussed under three categories in roots of plants inoculated with T22 under untreated control conditions relative to alkali-exposed plants inoculated with T22 (Fig. 3D; Supplementary Table S3).

We also analyzed the biological significance of differentially expressed genes (DEGs) in the roots of alkali-exposed Sugar Snap inoculated with T22 using the ShinyGO (v. 0.80) bioinformatics tool. The pathways significantly enriched in the upregulated DEGs were associated with various biological processes (response to stimulus, organic acid catabolic process, L-phenylalanine catabolic process), molecular functions (ammonia-lyase activity, heme binding), and cellular components (intracellular organelles, organelles) in plants (Fig. 4A-4C). Additionally, several KEGG pathways were enriched among the significantly upregulated DEGs (flavonoid biosynthesis, biosynthesis of secondary metabolites, phenylalanine metabolism, phenylpropanoid biosynthesis, isoflavonoid biosynthesis) and downregulated DEGs (plant-pathogen interaction, MAPK signaling pathway-plant, plant hormone signal transduction, ribosome) in Fe-deficient plants inoculated with T22 (Fig. 4D). We further analyzed the co-expressed network DEGs using WGCNA, which revealed several KEGG pathways with varying degrees of enrichment between Fe-T22 and Fe-conditions (Fig. 4E). We found that phenylpropanoid biosynthesis, MAPK signaling pathway, flavonoid biosynthesis, and isoflavonoid biosynthesis were among the top significantly upregulated pathways (green nodes). In contrast, pathways associated with ribosome, oxidative phosphorylation, DNA replication, terpenoid backbone biosynthesis, and nucleotide excision repair were among the most significantly downregulated pathways (red nodes) under these conditions (Fig. 4E).

**Fig. 4.**
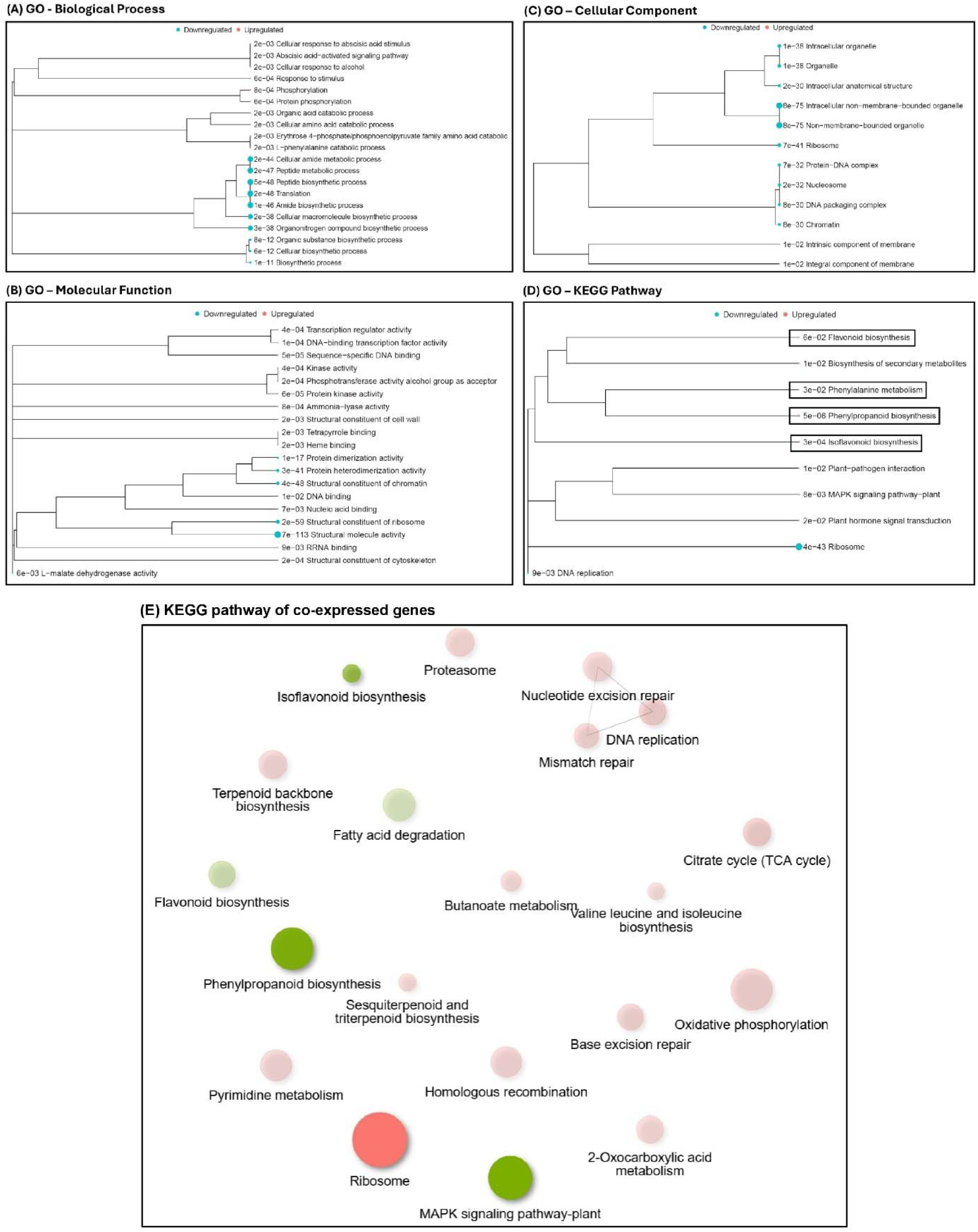
Gene enrichment and functional analysis of differentially expressed genes (DEGs) in pea roots cultivated under alkaline stress inoculated with T22: (A) biological process, (B) molecular function, (C) cellular component (D) KEGG pathway analysis and (E) co-expression network analysis of KEGG pathways for DEGs between alkaline+T22 and alkaline conditions. The analysis was performed using ShinyGO 0.76.3, with separate categories for differentially (significance cutoff: *FDR < 0.1*)) upregulated and downregulated genes indicated by blue and red dots, respectively. For using WGCNA (weighted gene co-expression network analysis), green and red represent up- and down-regulated pathways, respectively. Bigger nodes represent larger gene sets. Thicker edges represent more overlapped genes.

### 3.6 Effect of cinnamic acid on the response of alkali-exposed pea plants

Given the RNA-seq analysis indicating the involvement of genes linked to the flavonoid biosynthetic pathway in T22-mediated mitigation of alkaline stress, we conducted a targeted study to investigate whether cinnamic acid (CA) could elevate flavonoid levels and induce alkaline stress tolerance in pea plants without exogenous T22. We observed that alkali-exposed plants exhibited significantly lower SPAD scores and Fv/Fm ratios compared to controls. However, plants treated with either cinnamic acid or T22, regardless of soil pH, showed SPAD scores and Fv/Fm ratios statistically similar to those of control plants (Fig. 5A-5B). Plant height and dry weight (both root and shoot) also declined under alkaline stress. Nevertheless, alkali-stressed plants supplemented with either T22 or cinnamic acid displayed a significant increase in plant height and dry weight compared to alkali-imposed plants without any treatment. Plants treated with T22 or cinnamic acid under control conditions showed similar plant height and dry weight to controls (Fig. 5C-5D). Overall, the findings of this study align with the observed upregulation of genes within the flavonoid biosynthetic pathway in the RNA-Seq results.

**Fig. 5.**
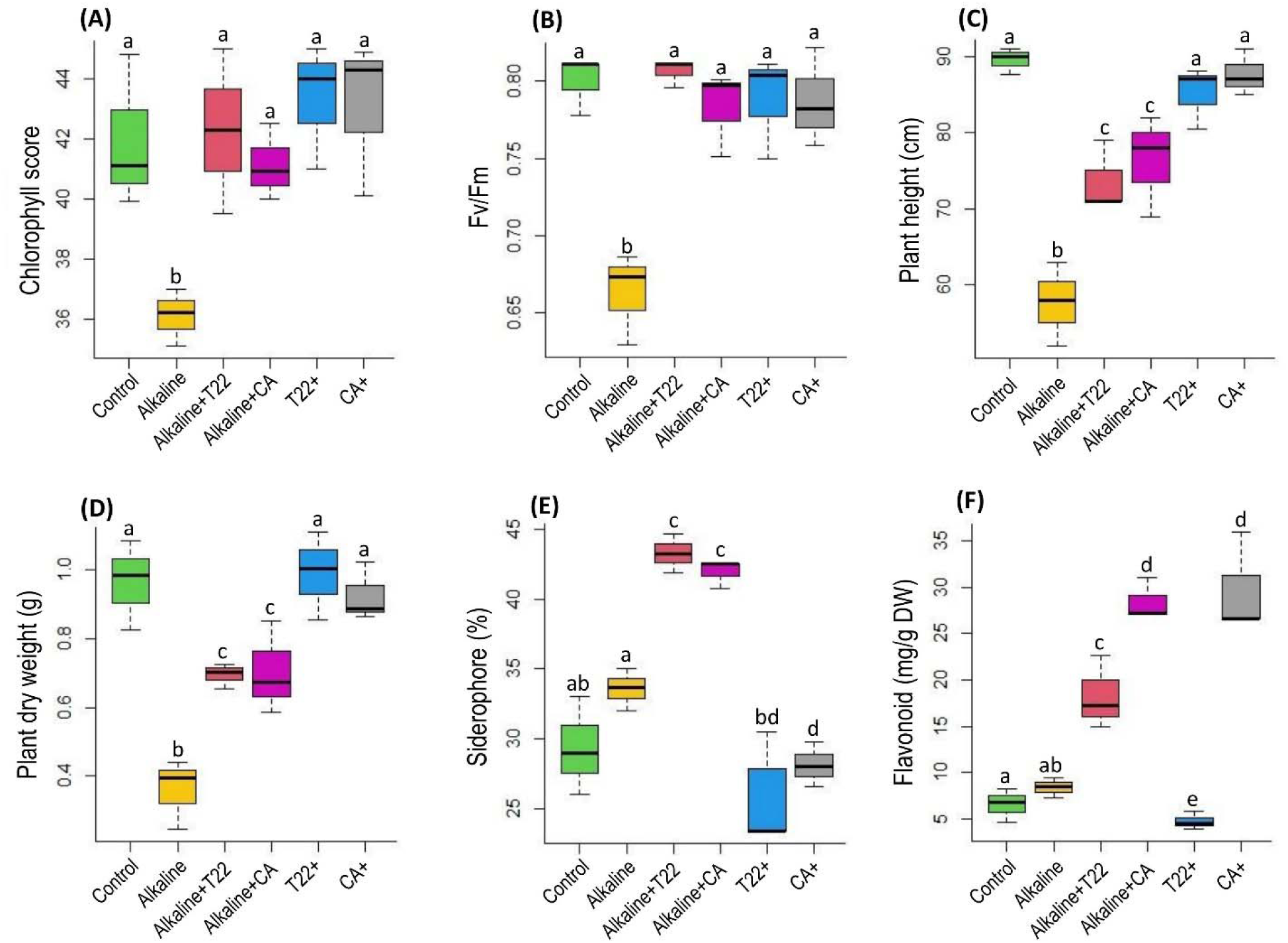
Effect of cinnamic acid (CA) on leaf SPAD score (A), Fv/Fm: maximal quantum yield of PS II (B), plant height (C), plant dry weight (D), rhizosphere siderophore (E) and root flavonoid content (F) in pea cultivated with or without the absence (control) or presence of alkaline stress and T22. The data presented are means with standard deviations (*n* = 3 individual replicates). Statistically significant differences in the ANOVA test among different treatments (*p* < 0.05) are indicated by different letters above the bars.

Siderophore levels in the rhizosphere soil showed no significant changes under alkaline stress compared to controls (Fig. 5E). Alkali-exposed plants supplemented with T22 or cinnamic acid showed a significant increase in siderophore levels compared to plants solely cultivated with soil alkalinity. Siderophore levels in plants without soil alkalinity, treated with either T22 or cinnamic acid, were similar to the control plants (Fig. 5E). Furthermore, total flavonoid content in roots showed no significant changes under alkaline stress relative to controls. However, alkali-imposed plants supplemented with T22 or cinnamic acid exhibited a significant increase in flavonoid content compared to alkali-stressed plants (Fig. 5F). Interestingly, control plants inoculated with T22 had significantly lower flavonoid levels compared to all other treatment groups. In contrast, alkali-exposed plants treated with cinnamic acid showed significantly higher flavonoid levels than those of controls (Fig. 5F).

### 3.7 Changes in nodulation and rhizobia dynamics in roots nodules

The number of root nodules significantly declined under alkaline stress compared to the control. However, the addition of T22 with or without soil alkalinity led to a significant increase in the number of root nodules, with levels returning to those similar to the controls (Fig. 6A). Principal component analysis (PCA) revealed substantial variation (88.4%), resulting in distinct separation of sample groups (Fig. 6B). We also analyzed species richness, diversity, and relative abundance of 16S bacterial communities in root nodules. Species richness (Chao1) and species diversity (Shannon index) showed no significant variations in bacterial communities in the root nodules of Sugar Snap in the presence or absence of alkaline stress and T22 inoculation (Fig. 6C-6D).

**Fig. 6.**
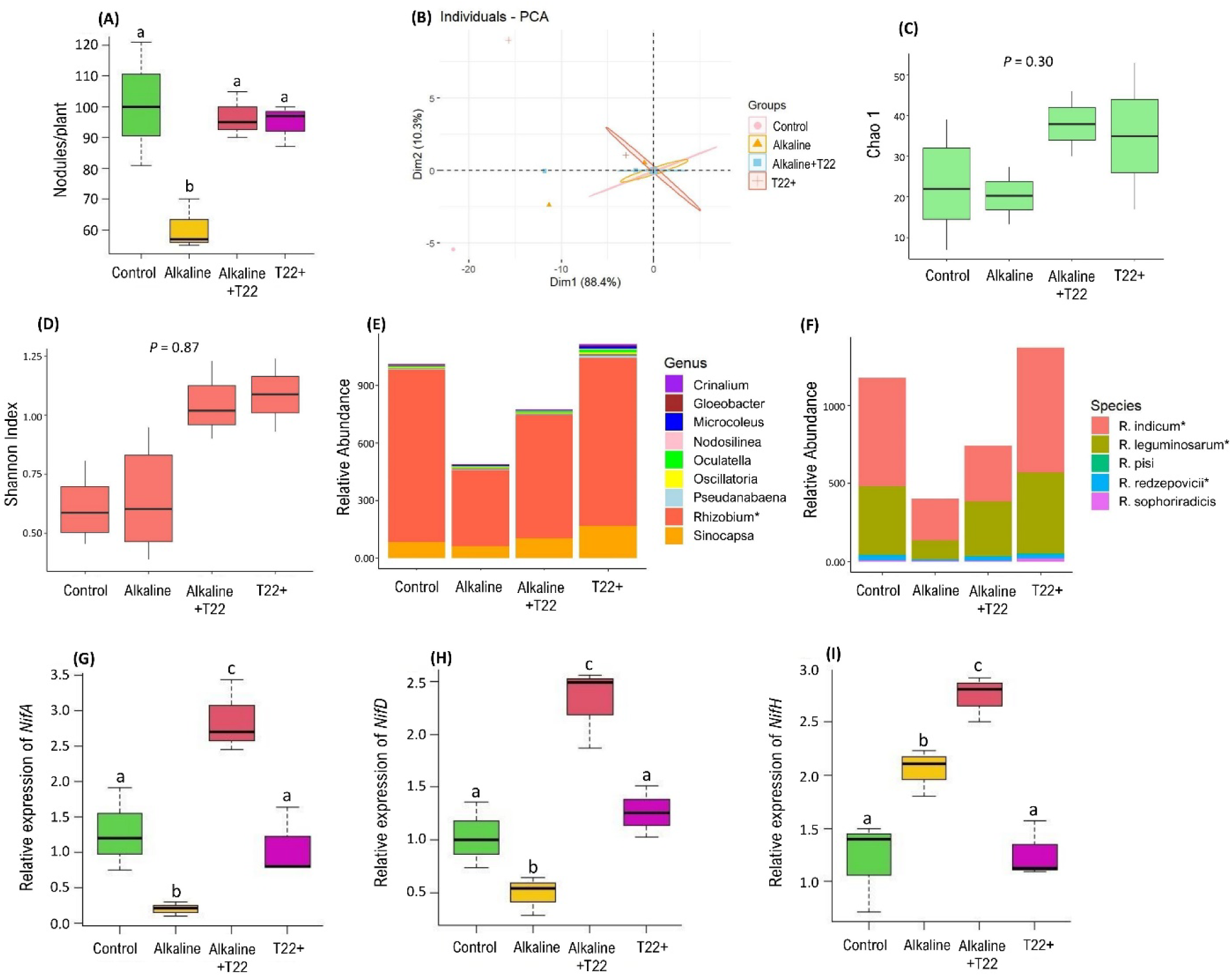
Changes in nodule number (A), principal component analysis (B), Chao 1 (C), Shannon index (D), relative abundance of bacterial genera (E), relative abundance of *Rhizobium* species (F) as well as the expression of *NifA* (G), *NifD* (H) and *NifH* (I) in nodules of pea plants inoculated with or without T22 cultivated in the absence or presence of alkaline stress. The data presented are means with standard deviations (*n* = 3 individual replicates). Statistically significant differences in the ANOVA test among different treatments (*p* < 0.05) are indicated by different letters above the bars, where applicable.

Nanopore sequencing revealed the differential relative abundance of bacterial genera, with a notable presence of the *Rhizobium* genus in root nodules influenced by alkaline stress and T22 inoculation (Fig. 6E). In the 16S analysis, the relative abundance of *Rhizobium* significantly decreased in nodules under alkaline stress but increased significantly when T22 was supplemented in alkaline soil. Plants inoculated with T22 without soil alkalinity showed a relative abundance of *Rhizobium* similar to that of the controls (Fig. 6E). Additionally, *Sinocapsa* and *Crinalium* did not show substantial changes in root nodules between the control and alkaline soil, regardless of T22 inoculation. However, plants inoculated with T22 under control conditions exhibited a slight increase in the relative abundance of *Sinocapsa* and *Crinalium* compared to other treatment groups (Fig. 6E). We observed no changes in the abundance of *Gloeobacter*, *Microcoleus*, *Nodosilinea*, *Oculatella*, *Oscillatoria*, and *Pseudanabaena* in nodules, regardless of soil pH or T22 inoculation (Fig. 6E). Given that *Rhizobium* species are crucial for legume nodulation, we further analyzed at the species level. The relative abundance of *R. indicum*, *R. leguminosarum*, and *R. redzepovicii* significantly decreased in root nodules under alkaline stress compared to controls (Fig. 6F). However, alkali-stressed plants inoculated with T22 showed a significant increase in the relative abundance of *R. leguminosarum* and *R. redzepovicii* compared to alkaline conditions. Plants inoculated with T22 without soil alkalinity exhibited a similar relative abundance of *R. indicum*, *R. leguminosarum*, and *R. redzepovicii* in root nodules to controls (Fig. 6F).

In this study, alkaline stress caused a significant decrease in the expression of *NifA* and *NifD* in nodules compared to controls (Fig. 6G-6H). However, alkali-imposed plants inoculated with T22 showed a significant increase in *NifA* and *NifD* expression compared to alkali-exposed plants without T22. Control plants supplemented with T22 exhibited similar levels of *NifA* and *NifD* expression in nodules to those of controls (Fig. 6G-6H). Furthermore, the expression of *NifH* significantly increased under alkaline stress compared to controls (Fig. 6I). The addition of T22 to alkali-exposed plants resulted in a significant increase in *NifH* expression in nodules relative to alkali-stressed plants without T22. Control plants inoculated with T22 showed a similar *NifH* expression pattern to that of controls (Fig. 6I).

### 3.8 Changes in plant responses supplemented with inorganic Fe

We cultivated Sugar Snap for 5 weeks to compare the effects of alkaline stress with T22 and FeEDDHA supplementation. We observed that leaf SPAD scores significantly declined under alkaline stress compared to controls. However, the addition of either FeEDDHA or T22 inoculum to the soil resulted in a significant increase in SPAD scores compared to alkali-exposed plants (Fig. 7A). T22 inoculated without soil alkalinity exhibited a similar SPAD score to that of controls and alkali-exposed plants supplemented with FeEDDHA. Notably, control plants supplemented with FeEDDHA showed a significantly higher SPAD score than all other treatment groups (Fig. 7A). In terms of plant height and biomass, alkaline stress caused a significant reduction compared to controls. The addition of T22 or FeEDDHA significantly improved these morphological traits relative to plants grown solely under soil alkalinity (Fig. 7B-7C). Plants without soil alkalinity supplemented with either T22 or FeEDDHA exhibited similar plant height and biomass to controls and alkali-exposed plants supplemented with FeEDDHA (Fig. 7B-7C). Interestingly, plants exhibited poor root development under alkaline stress compared to the control (Supplementary Fig. S6). In contrast, the alkaline+T22 treatment significantly improved root development, whereas FeEDDHA+ did not show any improvement under soil alkalinity. Both T22+ and FeEDDHA+ treatments under control conditions showed root development comparable to the control (Supplementary Fig. S6).

**Fig. 7.**
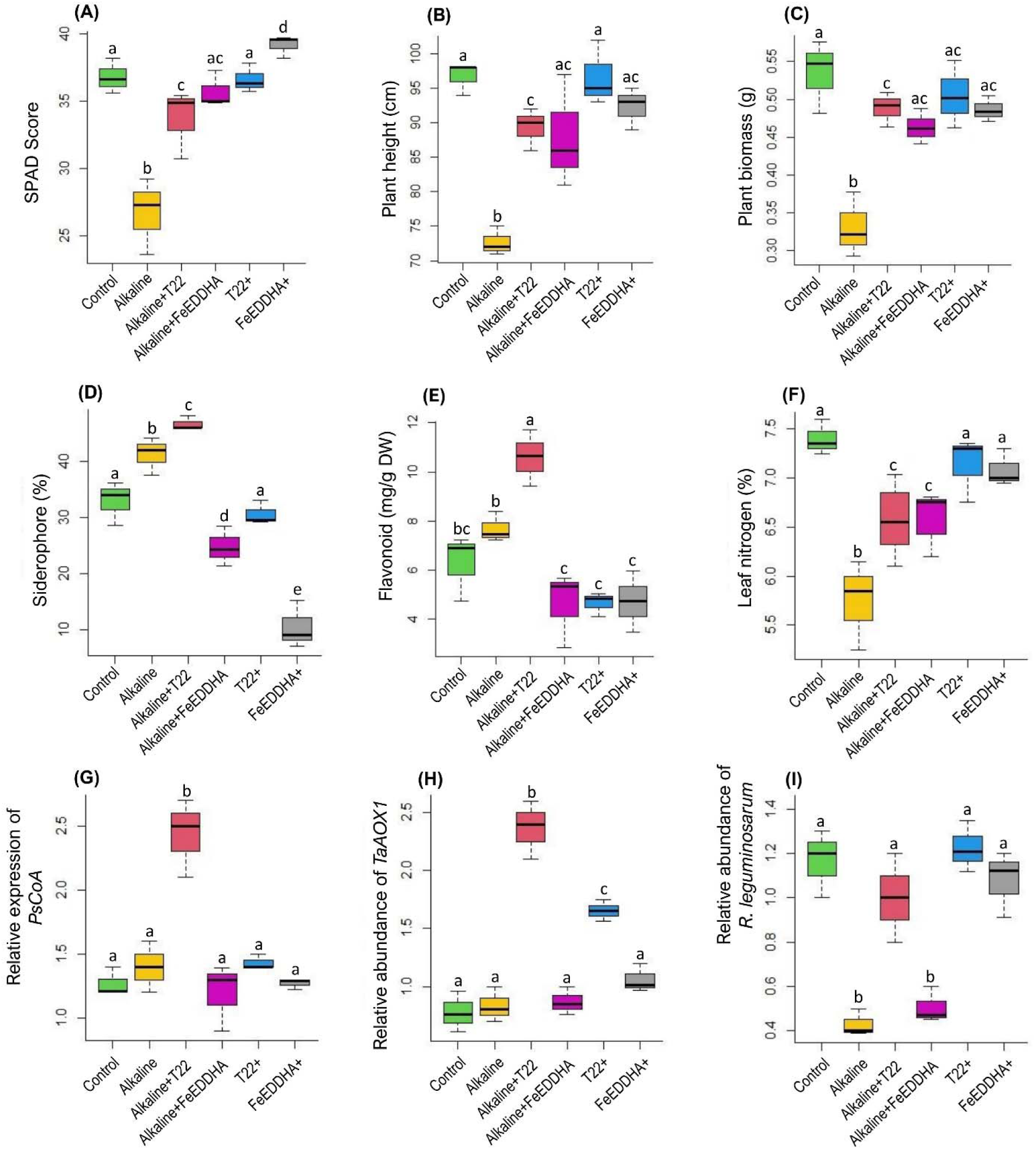
Effect of FeEDDHA on leaf SPAD score (A), plant height (B), plant dry biomass (C), rhizosphere siderophore (D), root flavonoid content (E) and leaf nitrogen (F), relative expression of *PsCoA* in roots (G), relative abundance of *TaAOX1* (H) and *R. leguminosarum* (I) in nodules of pea plants cultivated with or without the absence (control) or presence of alkaline stress and T22. The data presented are means with standard deviations (*n* = 3 individual replicates). Statistically significant differences in the ANOVA test among different treatments (*p* < 0.05) are indicated by different letters above the bars.

Analysis of rhizosphere siderophore levels revealed a significant increase under alkaline stress compared to controls (Fig. 7D). However, alkali-exposed plants inoculated with T22 showed a significant increase in siderophore levels compared to those grown solely under alkaline stress. In contrast, alkali-exposed plants supplemented with FeEDDHA exhibited a marked decrease in siderophore levels compared to controls, Fe-deficient plants, and alkali-imposed plants inoculated with T22. Control plants inoculated with T22 showed siderophore levels similar to those observed in controls. Furthermore, FeEDDHA supplementation in control conditions resulted in significantly lower siderophore levels compared to all other treatment groups (Fig. 7D).

In this study, the total flavonoid content in the roots did not show significant changes due to alkaline stress compared to controls. However, alkali-stressed plants inoculated with T22 exhibited a significant increase in flavonoid content relative to those grown solely in control or alkali-exposed soil (Fig. 7E). Interestingly, alkali-stressed plants treated with FeEDDHA and control plants supplemented with either T22 or FeEDDHA showed a significant decrease in flavonoid content compared to the other two treatment groups (Fe- and Fe-T22+) (Fig. 7E). Furthermore, leaf nitrogen content significantly decreased under alkaline stress compared to controls (Fig. 7F). Alkali-exposed plants treated with either T22 or FeEDDHA showed a significant increase in nitrogen content compared to alkali-stressed plants without these treatments. Finally, Fe-sufficient plants inoculated with T22 or supplemented with FeEDDHA exhibited similar leaf nitrogen content to controls. Interestingly, alkali-imposed plants supplemented with FeEDDHA showed similar nitrogen levels in the leaves to those of controls (Fig. 7F).

We further investigated the expression pattern of *PsCoA*, which showed a significant increase in the roots of alkali-stressed plants inoculated with T22 compared to controls and other treatment groups (Fe-, FeEDDHA, T22+, and FeEDDHA+) (Fig. 7G). The relative abundance of T22 (*TaAOX1*) in the roots significantly increased when T22 was introduced into the soil, both under alkaline and control conditions, relative to controls and FeEDDHA-treated soil. Particularly, the relative abundance of T22 was higher under alkaline stress compared to controls (Fig. 7H). Furthermore, soil alkalinity led to a significant reduction in the relative abundance of *R. leguminosarum* in nodules, with no significant changes observed with FeEDDHA supplementation (Fig. 7I). However, the addition of T22 with or without soil alkalinity and FeEDDHA addition under control, resulted in similar levels of *R. leguminosarum* in the nodules as observed in the controls (Fig. 7I).

We also investigated the interaction between T22 and *R. leguminosarum* under varying pH levels in nutrient media. In control media, the growth of both T22 and *R. leguminosarum* was significantly reduced when co-cultured compared to their growth when cultured alone (Supplementary Fig. S7). Under alkaline conditions, T22 growth remained unchanged whether cultured alone or co-cultured with *R. leguminosarum*. In contrast, the growth of *R. leguminosarum* significantly increased when co-cultured with T22 in alkali-imposed media compared to when it was cultured alone (Supplementary Fig. S7).

## 4 Discussion

Soil alkalinity and its adverse effects on mineral availability, particularly Fe, pose a major challenge in agriculture, frequently leading to stunted growth and reduced yields in legume crops (Bhat et al., 2024; Brear et al. 2013). Traditional approaches, such as genetic modification and chemical fertilizers, aimed at enhancing alkalinity tolerance may lack sustainability and long-term effectiveness (Yadav et al. 2023; Guzmán et al. 2022). A microbiome-based approach, specifically utilizing beneficial microorganisms like T22, offers a sustainable alternative to enhance crop resilience to alkali-mediated nutrient deficiency (Kabir et al. 2022; Zhao et al. 2020). This study provides the first mechanistic insights into how T22 establishes a symbiotic relationship with the host, modulates the root transcriptome, and reshapes nodule microbial dynamics to enhance alkalinity tolerance in garden peas.

### 4.1 The colonization efficiency of T22 in mitigating soil alkalinity is host-specific

Fungal colonization in plant roots is highly host-specific, influenced by factors such as host genetics, immune responses, and the compatibility of fungal species with the plant’s physiology (Morán-Diez et al. 2015; Liu et al. 2019). In this study, the abundance analysis of the *TaAOX1* gene indicated that only the Sugar Bon and Sugar Snap cultivars exhibited significant fungal colonization. This suggests a genetic basis for host-specific interactions, which could be utilized to enhance fungal symbiosis and stress tolerance in cultivars that are more efficient with microbiomes. Morán-Diez et al. (2015) analyzed the transcriptomic responses of *Trichoderma virens* interacting with maize and tomato roots, identifying 35 genes with differential expression between the two hosts, including 10 genes with higher expression in tomato roots. In this study, Sugar Snap inoculated with T22 exhibited the highest fungal colonization and enhanced photosynthetic attributes under alkaline stress, whereas colonization significantly decreased when mineral availability was not disrupted by soil alkalinity. This highlights the dependency of T22’s beneficial effects on the plant’s mineral status, particularly Fe, and implies the complex interplay between the fungus, the plant, and soil nutrient availability.

Furthermore, partitioning the Sugar Snap root system revealed the highest T22 colonization in the meristematic zone, as indicated by the abundance of *TaAOX1*. The meristematic zone offers rich nutrients, exudates, and a favorable microenvironment, enhancing microbial colonization compared to other root segments (Tsai et al. 2023; Canarini et al. 2019). The role of the meristematic zone of roots in water and mineral absorption and its variation in fungal colonization across root segments have been reported in other studies (Kabir and Bennetzen 2024; López-Bucio et al. 2003). The high T22 abundance in the meristematic zone may be linked to active cell division and growth, providing optimal conditions for this fungal species (Russo et al. 2019). Root partition studies are essential for understanding microbial colonization and interactions, offering insights into how different root sections influence and are influenced by microbial communities (Bart et al. 2022; Habiyaremye et al. 2020). These findings suggest the importance of screening cultivars for favorable T22 colonization to optimize plant-microbe interactions and enhance stress resilience in crops.

### 4.2 Host flavonoid-mediated modulation of factors promoting symbiosis

Microbial networks facilitate the exchange of signals and cues among plants, triggering adaptive responses to environmental stress (Gorzelak et al. 2015). Symbiotic signaling between microbes and plants exposed to abiotic stress is essential for mediating beneficial interactions that improve plant fitness and resilience. In this study, a split-root assay revealed that the presence of T22 in the rhizosphere, even when split into two distinct compartments, is not necessary to induce alkaline stress tolerance in garden pea. We found that resilience to alkaline stress in Sugar Snap was maintained even when T22 was inoculated in only one root compartment, suggesting that the beneficial effects of T22 extend beyond localized root interactions. This suggests that the mechanism through which T22 induces alkaline stress tolerance in garden pea relies on the whole plant interactions triggered by the host plant itself, rather than the root-borne associations within the rhizosphere. However, the determinants that aid colonization are influenced by host genetics, which is typically linked to the secretion of metabolites and responses to environmental cues (Wiesmann et al. 2023; Bapaume and Reinhardt 2012).

Our RNA-seq analysis of the meristematic segment of roots revealed the upregulation of several key genes associated with symbiosis in alkali-exposed pea plants inoculated with T22. Particularly, we observed the induction of genes responsible for monooxygenase activity, ammonia-lyase activity, auxin-responsive proteins, and 4-coumarate A ligase in alkali-stressed plants inoculated with T22. Monooxygenase enzymes are crucial for maintaining symbiotic relationships between plants and microorganisms, playing roles in signaling, secondary metabolite production, and hormone regulation that enhance plant health and resilience in symbiotic systems (Wu et al. 2020; Mnguni et al. 2020; Dasgupta et al. 2011). Similarly, ammonia-lyase enzymes, particularly phenylalanine ammonia-lyase (PAL), are significant in plant-microbe symbioses. PAL is integral to the phenylpropanoid pathway, converting phenylalanine to cinnamic acid, which is then transformed into 4-coumarate - a key component in the biosynthesis of coumarins and flavonoids (Li et al. 2020). Interestingly, the co-expressed gene network consistently showed enrichment in pathways related to flavonoid biosynthesis, including flavonoid, isoflavonoid, and phenylpropanoid biosynthesis, reinforcing the role of flavonoids in pea plants inoculated with T22 under alkaline stress. Coumarins, synthesized through the phenylpropanoid pathway, act as signaling molecules that regulate symbiotic interactions between microbes and plants (Harbort et al. 2020; Stringlis et al. 2009). In Fe-deficient *Arabidopsis*, 4-coumarate is crucial for enhancing Fe uptake, which coordinates coumarin biosynthesis to optimize Fe acquisition (Paffrath et al. 2024). Coumarins can also modify the root microbiota, addressing Fe deficiency in *Arabidopsis* (Harbort et al. 2020). Furthermore, the accumulation of flavonoids in plants is controlled by genes within the flavonoid biosynthetic pathway (Borevitz et al. 2000). Flavonoids released by plants promote microbial colonization and act as pre-symbiotic signaling molecules (Lidoy et al. 2023; Schliemann et al. 2008). Consistent with these findings, KEGG pathway analysis of T22-inoculated roots showed enrichment in flavonoid, isoflavonoid, and phenylpropanoid biosynthetic pathways. Flavonoids are crucial for rhizobia nodulation in legumes, acting as key signaling molecules that attract rhizobia to the root and stimulate bacterial gene expression. NodD proteins in rhizobia specifically recognize and respond to flavonoid signals secreted by legumes, leading to nodulation initiation (Wang et al. 2018; Peck et al. 2006). Additionally, flavonoids serve as antioxidants, protecting plants against environmental stress (Grace 2000). Thus, our findings suggest that T22-mediated induction of ammonia-lyase and 4-coumarate A ligase in pea roots may be crucial for the initial steps of flavonoid biosynthesis. This could lead to the formation of various flavonoid compounds that promote symbiotic associations with T22 and/or rhizobia, helping pea plants cope with alkali-exposed nutrient deficiency.

To further validate the role of flavonoids in potential symbiotic relationships with soil microbes and their contribution to alkaline stress tolerance in pea plants, we conducted a targeted study on the effect of cinnamic acid, a precursor in the flavonoid biosynthesis pathway. Our study showed that pea plants either inoculated with T22 or supplemented with cinnamic acid exhibited significant improvements in photosynthetic parameters and plant biomass compared to alkali-exposed plants. This enhancement in plant health was associated with increased synthesis of rhizosphere siderophores and flavonoids in the roots. Since siderophores are essential for Fe solubilization and uptake, the increased siderophore levels in T22-inoculated plants suggest that T22 enhances Fe acquisition by stimulating microbial activity in the rhizosphere. Previous studies have shown that siderophore-producing microbes improve Fe uptake efficiency in alkali-mediated Fe-deficient soils (Jin et al., 2014; Kabir et al., 2022). Furthermore, elevated flavonoid levels may stimulate the release of siderophores by rhizosphere microbes, thereby promoting the solubilization of insoluble Fe and aiding pea plants in coping with alkaline stress. Zamboni et al. (2012) found that tomato plants respond to Fe deficiency by upregulating the flavonoid-3-hydroxylase gene, indicating a role for flavonoids in influencing Fe bioavailability in soil. In another study, *T. harzianum* application increased flavonoid content in tomato plants, leading to enhanced photosynthetic characteristics and improved fruit quality (Vukelić et al. 2021). Furthermore, microbial consortia containing *T. afroharzianum* 5F have been shown to enhance flavonoid and phenol levels in bean plants infected with *Rhizoctonia bataticola* (Singh et al., 2024). Thus, it appears that flavonoids may play diverse roles in regulating both plant development and interactions with commensal and pathogenic microbes (Wang et al. 2022). Our findings align with these reports, indicating that T22 not only improves plant Fe status but also stimulates beneficial microbial activity in the rhizosphere of pea exposed to alkaline stress. Future research should investigate whether additional flavonoid derivatives or microbial metabolites play a role in mineral homeostasis and plant-microbe interactions, potentially identifying new pathways for enhancing crop resilience.

### 4.3 Reversal of alkali-induced retardation in mineral and redox status

Once a symbiotic relationship between a plant and a microbe is established, host plants often reprogram their responses to better cope with abiotic stress (Kabir and Bennetzen 2024; Meena et al. 2017). In this study, garden pea plants cultivated under soil alkalinity exhibited reduced SPAD scores and decreased photosynthetic efficiency, aligning with previous findings on legume crops under alkali-mediated Fe shortage (Haque et al. 2022; Kabir et al. 2012). In contrast, inoculation with T22 significantly improved chlorophyll synthesis and photosynthetic performance in these alkali-imposed plants. This improvement is consistent with earlier studies reporting enhanced SPAD scores, photosystem II activity, and photosynthesis performance index following T22 application to mineral-deficient plants (Kabir and Bennetzen 2024; Kabir et al. 2022; Singh et al. 2019).

Soil alkalinity causes essential micronutrients to precipitate or become chemically bound in forms that plants cannot absorb (Dhaliwal et al. 2019). We also observed a decrease in Fe and Mn levels in garden pea plants exposed to alkaline stress. However, inoculation with T22 restored Fe and Mn levels in both roots and shoots. Previous research showed that *Trichoderma* strains can elevate Fe levels but may reduce concentrations of Cu, Mn, and Zn in some plants grown in calcareous soils (Kabir and Bennetzen 2024; Santiago et al. 2011). These differential mineral levels may result from variations in mineral forms and changes in metal transporter activity in host plants (Ye et al. 2017). Also, several micronutrient transporters exhibit overlapping specificity, facilitating the uptake of multiple microelements in plants (Rai et al. 2021). In this study, the enhancement of Fe and Mn status in alkali-exposed plants due to T22 implies its potential role in inducing key transporters for nutrient acquisition, as evidenced by the upregulation of specific genes in roots. Therefore, the recovery of Fe may be mediated by siderophores released by T22, while the improvement in Mn levels could be a secondary effect of the symbiotic benefits conferred by T22 under soil alkalinity. Consistently, RNA-seq analysis identified genes related to manganese ion binding, Fe ion binding, and metal ion binding that were either downregulated or showed no change but were substantially induced by T22 under alkaline stress. The variability in resistance to soil alkalinity among different plant cultivars is influenced by distinct mechanisms of homeostasis and gene expression patterns (Atencio et al. 2021; Kabir et al. 2012). In our study, alkali-exposed pea plants inoculated with T22 exhibited upregulation of Fe-related genes (*Psat3g159880*, *Psat5g286200*) compared to non-inoculated plants, suggesting a symbiotic advantage in enhancing Fe uptake. This indicates that alkali-exposed plants may either sense a need for increased Fe uptake or are driven to induce Fe uptake when insoluble Fe(III) is solubilized through microbial interactions in the rhizosphere. Elevated siderophore levels in the rhizosphere under soil alkalinity, especially when T22 was supplemented, support this hypothesis. Siderophores are typically released by bacterial and fungal communities to chelate Fe in Fe-deficient conditions (Jin et al. 2010; Crowley et al. 2006). Harzianic acid, a secondary metabolite produced by T22, binds with Fe(III) and enhances Fe solubilization in the soil, benefiting both other microorganisms and the host plant (Vinale et al. 2013). The variability in siderophore levels released by *Trichoderma*, influenced by host cultivar implies that elevated siderophore levels in the alkali-supplemented rhizosphere are closely associated with exogenous T22, particularly in the roots of Sugar Snap.

The accumulation of reactive oxygen species (ROS) due to mineral deficiency leads to oxidative damage in cellular components, including lipids, proteins, and DNA (Tewari et al. 2013). Since Fe is crucial for primary antioxidant enzymes, oxidative stress is exacerbated in plants under mineral deficiency (Halliwell 2006). In our study, alkali-exposed pea plants inoculated with T22 not only exhibited upregulation of mineral transporters but also showed increased expression of peroxidase and glutathione-related genes. Consistently, the biochemical analysis of H_2_O_2_ also showed a significant decline in the roots of alkali-stressed plants inoculated with T22. This suggests that the symbiotic relationship with T22 might enhance ROS scavenging in plants under alkaline stress. Although the precise mechanisms by which ROS balance is regulated in plants with mycorrhizal associations are still not fully understood, several studies have highlighted the role of AMF and *Trichoderma* in inducing antioxidants to protect plants from abiotic stressors, including Fe deficiency (García-Sanchez et al. 2014; Kabir and Bennetzen 2024). Flavonoids, known for their antioxidant properties, play a crucial role in detoxifying and scavenging ROS produced as by-products of oxidative metabolism (Jan et al. 2021; Nakabayashi et al. 2014). For instance, Mastouri et al. (2012) demonstrated that T22 colonization in drought-stressed tomato plants enhanced ROS scavenging capabilities and the recycling of oxidized ascorbate and glutathione, contributing to improved ROS-scavenging pathways. Additionally, T22 has been shown to boost antioxidant defense in alkali-mediated Fe-deficient soybeans by increasing glutathione reductase activity and levels of glutathione and methionine, which stabilize the redox state (Kabir et al. 2022). Although the specific mechanisms of T22 playing a role in ROS scavenging require further investigation, our findings suggest that T22-mediated increases in flavonoid levels may positively influence ROS homeostasis in alkali-exposed pea plants. Furthermore, maintaining ROS balance can improve photosystem II efficiency, ensuring better light capture and energy conversion in plants exposed to abiotic stress (Nadarajah 2020), and support meristem development by minimizing cellular damage (Huang et al. 2019). Excessive buildup of ROS, especially in photosystems I and II, can lead to damage of Fe-S proteins, essential cofactors in plant photosynthesis (Xiong et al. 2021; Murchie and Niyogi 2011). Our findings indicate that alkaline stress resulted in reduced chlorophyll synthesis and decreased Fv/Fm in the leaves of garden pea. Since alkali-stressed pea inoculated with T22 substantially improved mineral and ROS status, genes related to photosystem II (*Psat1g002320*) and meristem development (*Psat5g186720*) were consistently upregulated under these conditions. This molecular evidence was supported by improved leaf Fv/Fm ratios and root systems in plants supplemented with T22 under alkaline stress. Research has shown that mycorrhizal symbiosis mediated by AMF in *Robinia pseudoacacia* enhances PSII activity in chloroplasts, leading to improved water use efficiency (Yang et al. 2015). Additionally, studies have shown that T22 can restore chlorophyll levels and photosystem II activity in other mineral-deficient plants (Kabir et al. 2022; Singh et al. 2019). Our study suggests that T22 may play a crucial role in enhancing photosynthetic efficiency, potentially by indirectly modulating ROS homeostasis.

### 4.4 T22 restores nodule bacterial dynamics in pea exposed to alkaline stress

Rhizobia is essential for the growth and stress resilience of legume crops, as they form symbiotic relationships that enhance nitrogen fixation and improve overall plant health under various stress conditions (Shumilina et al. 2023). Given the observed changes in nodulation and expression of nitrogen-fixation genes, we further investigated whether T22 impacts the bacterial diversity and dynamics in root nodules of pea plants exposed to alkaline stress. Our results demonstrate that soil alkalinity leads to a marked reduction in the number of root nodules. However, the addition of T22 not only mitigated the negative effects of alkaline stress but also enhanced nodule formation regardless of the pH levels of the soil. This suggests a potential biostimulant effect of T22 on nodule development. The synergistic effects of *Rhizobium* and mycorrhizal interactions are still not fully understood. However, rhizobia and AMF can form symbioses with legumes, significantly enhancing plant mineral nutrition, particularly nitrogen and phosphorus uptake (Ossler et al. 2015; Xie et al. 2018). In this study, microbiome analysis in pea nodules showed substantial changes in species richness and diversity in response to pH levels and T22 inoculation. These findings highlight the profound impact of both soil alkalinity and T22 inoculation on the overall bacterial community structure within root nodules in pea plants. Particularly, Fe is essential for the functioning of nitrogenase, the enzyme complex responsible for nitrogen fixation in legume plant nodules (Schwember et al. 2019; Brear et al. 2013). In this study, the inoculation of alkali-exposed pea plants with T22 significantly increased the relative abundance of *Rhizobium*. In the co-culture experiment, *R. leguminosarum* colony size significantly increased due to T22 only under alkaline conditions, suggesting a positive interaction in which T22 complements *R. leguminosarum* growth in a condition-dependent manner. However, T22 itself does not exhibit enhanced growth in co-culture, implying a unidirectional benefit favoring *R. leguminosarum*. These findings suggest the positive impact of T22 on restoring beneficial *Rhizobium* populations under alkaline stress, thereby promoting nodulation and overall plant health of peas exposed to soil alkalinity. However, it is unknown at this point whether the improved nodule formation is a direct consequence of T22 on Rhizobia or the effect of the improved nutrient status of alkali-exposed pea plants indirectly mediated by T22. However, this study may encourage future research to identify the metabolites that drive beneficial microbe-microbe exchanges in high pH environments. Although Rhizobia-fungi symbiotic associations share common signals and processes, mycorrhizal symbiosis has been reported to increase phosphorus content. This can boost nitrogenase enzyme activity, leading to higher nitrogen fixation by rhizobial symbionts, and, in return, better mycorrhizal development (Ren et al. 2016; Hao et al. 2019). Additionally, AMF can reduce oxidative stress in the nodules (Porcel et al. 2003) and enhance the carbon metabolism within them (Goicoechea et al. 2004). Furthermore, fungi can produce and modulate plant hormones, which promote root and nodule development (Chanclud and Morel 2016). Recent advances in multi-omics technologies, such as metagenomics, can be leveraged to enhance our understanding of cues between microbial partners, potentially aiding the host plant in combating extreme environments (Han et al., 2024).

We further conducted a targeted study to elucidate whether T22-mediated mitigation of alkaline stress in pea plants is directly related to associations with rhizosphere microbes, including rhizobia, or indirectly linked to the improved Fe status of the plants. Fe fertilizers are commonly used to prevent the precipitation of Fe in the soil, ensuring its availability even in alkaline or calcareous soils, leading to healthier, more vigorous plant growth and increased yields (Schenkeveld et al. 2010). In this study, we found that FeEDDHA was as effective as T22 in mitigating the adverse effects of alkaline stress in chlorophyll content, plant height, and biomass caused by iron deficiency in pea plants. Furthermore, the addition of FeEDDHA under alkaline conditions resulted in no significant differences in these parameters compared to the non-treated controls. However, the non-significant differences in these parameters might be attributed to the reduced availability of other nutrients, such as Mn, or to the plant’s reduced ability to absorb FeEDDHA effectively due to root damage caused by soil alkalinity. Alkaline soils can hinder root development, which in turn limits the efficient uptake of essential minerals by plants (Saleem et al. 2023). Although FeEDDHA is stable in alkaline soil, some Fe may still bind to soil particles, limiting its bioavailability and effectiveness (Zuluaga et al. 2023). However, the improved root structure induced by T22 likely underlies more effective symbiotic interactions with beneficial soil microbes. *Trichoderma* is known to enhance root development through auxin-related signaling pathways (Contreras-Cornejo et al. 2024; Contreras-Cornejo et al. 2009).

Interestingly, we observed differences in rhizosphere siderophore levels, root flavonoid content, the expression of *PsCoA*, and the relative abundance of *TaAOX1* and *R. leguminosarum* - all of which showed a simultaneous decrease when alkali-exposed pea plants were supplemented with FeEDDHA compared to plants inoculated with T22. This suggests that the elevation of flavonoids and the restoration of *R. leguminosarum* largely depend on the biological interactions with T22 during the symbiotic process, rather than on the Fe status of the plants supplied through an exogenous source. Specifically, it implies that T22 was involved in siderophore production and host flavonoid content, potentially enhancing the plant’s nutrient acquisition processes, while FeEDDHA primarily provides a direct source of available Fe without stimulating these secondary metabolic pathways. Microbial siderophore production is closely linked to Fe availability in the soil (Timofeeva et al. 2022). When soil Fe levels are low, microorganisms increase siderophore production to scavenge Fe more effectively and make it available to both microbes and plants (Ahmed and Holmström 2014). Kabir and Bennetzen (2024) showed that lime-induced Fe deficiency significantly increased the siderophore-related (*SIT1*) gene in T22, resulting in the solubilization of Fe, which improved Fe status in sorghum. They found the highest siderophore accumulation in sorghum inoculated with T22 under alkaline conditions. Hence, the elevated siderophore released by T22 in the rhizosphere of Fe-deficient pea plants may be no longer needed when inorganic Fe fertilizer is available. However, the elevation of siderophores in the rhizosphere of alkali-exposed plants suggests that not only T22 but also other microbes may be involved. Several studies support the idea that bacterial communities, including rhizobia, release siderophores in response to soil alkalinity, thereby improving mineral uptake by plants alongside other beneficial interactions such as nitrogen fixation (Lurthy et al. 2020; Wang et al. 2024; Brear et al. 2013).

Our follow-up study further demonstrated that alkali-imposed pea plants were able to combat stress symptoms with Fe supplementation, without increasing rhizosphere siderophore and root flavonoid content, which is a requirement for T22-mediated improvement of plant health. The initiation of complex legume-*Rhizobium* development begins with the release of flavonoids (Dong and Song 2020). Since siderophores and flavonoids are involved in symbiotic associations, alkali-exposed plants bypassed this strategy by overcoming stress symptoms when inorganic Fe was supplemented in the soil. When nutrients are sufficient, bacteria may prioritize other metabolic processes over nodulation to conserve energy, which helps them efficiently allocate resources based on environmental cues, enhancing their overall fitness (Frawley and Fang 2014). Therefore, our findings suggest that the presence of T22, rather than Fe status alone, may be involved in restoring rhizobia dynamics under alkaline conditions, contributing to the mechanistic pathways that mitigate alkaline stress in pea plants. Thus, T22 may improve nodulation and support mineral uptake including Fe and Mn through specific mechanisms that go beyond simply altering the Fe status of the plants. This result is preliminary given the complexity of the soil system, which shapes multifactorial effects on alkali-exposed pea plants. So far, there is a lack of well-documented interaction between T22 and rhizobia in scientific literature. However, it is known that *Trichoderma* can enhance the activities of beneficial bacteria such as *Pseudomonas* or *Bacillus*, which contribute to improved plant health and nutrient uptake (Poveda et al. 2022; da Costa et al. 2022). *Trichoderma* species are known to produce a variety of metabolites that can play a role in shaping the composition and dynamics of soil or rhizosphere bacterial communities, influencing overall ecosystem health and plant-microbe interactions (Gao et al. 2023). This correlates with the levels of flavonoids in roots, as rhizobia are also attracted to flavonoid-mediated symbiosis (Shumilina et al. 2023; Li et al. 2019). Furthermore, *Trichoderma* application has been found to shift other fungal communities in the rhizosphere (Asghar and Kataoka 2021; Umadevi et al. 2018). Future research could explore how T22 interacts with rhizobia and identify the specific metabolites responsible for this association in legume plants under varying soil pH.

## 5 Conclusion

This study unveils the intricate mechanisms by which T22 interacts with both garden pea hosts and the rhizobia community to mitigate soil alkalinity. We demonstrated that T22 enhances plant health parameters under alkaline stress by improving tissue Fe and Mn levels and elevating rhizosphere siderophores. Furthermore, T22 fosters the 16S microbial community, particularly restoring the abundance of rhizobia and upregulating essential nitrogen fixation genes in nodules of pea exposed to alkaline stress (Fig. 8). Our findings also suggest that the beneficial effect of T22 in mitigating alkaline stress involves whole-plant interactions, not just the root system. RNA-seq analysis revealed significant gene expression changes linked to flavonoid biosynthesis, mineral transport, and redox homeostasis in T22-inoculated pea roots under soil alkalinity. Moreover, the restoration of plant health with cinnamic acid in the absence of T22 reinforces the role of flavonoids in promoting microbial symbiosis. However, the substitution of T22 with Fe supplementation reduced rhizosphere siderophore and flavonoid levels, as well as the abundance of *TaAOX1* and *R. leguminosarum*, nullifying the associations of microbiome-dominated improvement in plant health including Fe status. Overall, our mechanistic insights highlight the potential of T22 as a key player in microbiome-based strategies to alleviate alkaline stress in legume crops, paving the way for sustainable agricultural practices. These insights provide a valuable foundation for developing strategies to improve legume growth and stress resilience through microbial mining and targeted microbial interventions.

**Fig. 8.**
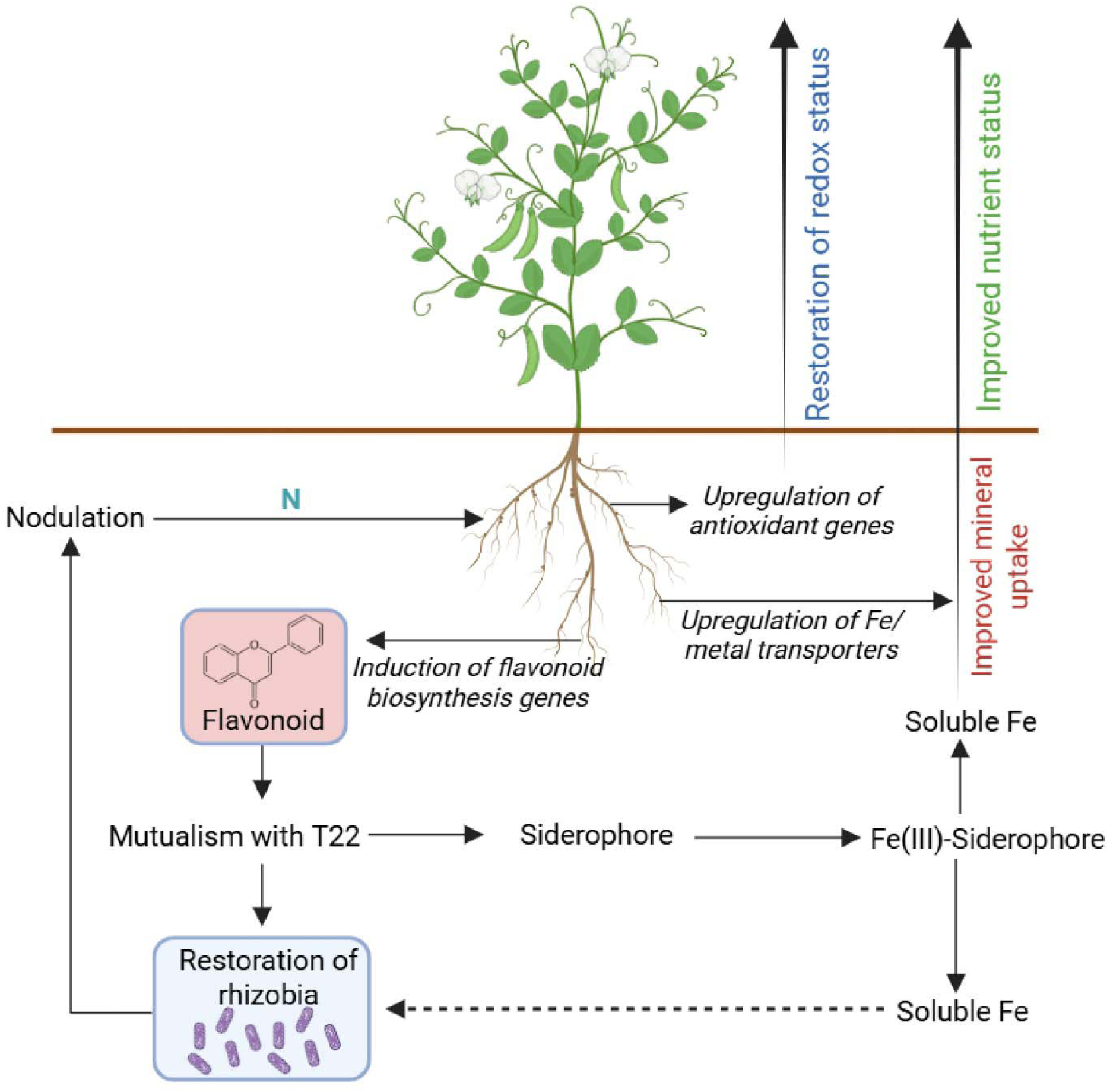
Model of interactions by which T22 influences the symbiotic processes to help pea plants cope with alkaline stress. T22 mainly enhances Fe availability through siderophore-mediated solubilization of Fe(III), increasing soluble Fe in the rhizosphere. The plants respond by upregulating Fe/metal transporters, leading to improved nutrient status. Additionally, T22 induces the expression of flavonoid biosynthesis genes, promoting flavonoid production, which facilitates mutualistic interactions with rhizobia and restores nodulation. Nodulation contributes to N assimilation, further supporting plant growth. Upregulation of antioxidant genes restores redox status, mitigating oxidative stress under alkaline stress. Overall, these interconnected pathways contribute to improved Fe homeostasis, nodulation efficiency, and alkaline stress resilience in pea plants inoculated with T22.

## Acknowledgments

We express our gratitude to the Genomics Core of Michigan State University. This research was supported by Louisiana Biomedical Research Network (grant ID: KAB004) and a startup grant (grant ID: 5SFAES-293007) from the University of Louisiana at Monroe.

## Author Contribution

AT was involved in experimental design, plant cultivation, RNA extraction, RNA-seq data analysis, data collection, interpretation, 16S data analysis, and prepared the manuscript draft. MRH assisted with DNA extraction and microbial co-culture experiments. AHK provided overall supervision, guidance in experimental design, data interpretation, and manuscript revision.

## Data Availability

The data supporting the findings of this study are available in the article and its supplementary information files. Also, the sequencing data used and described in this study has been uploaded to NCBI under the Bioproject accession numbers: PRJNA1133893 (RNA-seq) and PRJNA1134200 (16S).

## References

1. Ahmed, E. & Holmström, S. J. (2014) Siderophores in environmental research: roles and applications. Microbial Biotechnology, 7(3), 196–208.

2. Alcántara, E., Romera, F. J., Canete, M. & De la Guardia, M. D. (2000) Effects of bicarbonate and Fe supply on Fe (III) reducing capacity of roots and leaf chlorosis of the susceptible peach rootstock “Nemaguard”. Journal of Plant Nutrition, 23(11–12), 1607–1617.

3. Alexieva, V, Sergiev, Mapelli S, Karanov E. (2001) The effect of drought and ultraviolet radiation on growth and stress markers in pea and wheat. Plant Cell Environment 24, 1337– 1344.

4. Anders, S., Pyl, P.T. & Huber, W. (2015) HTSeq–a Python framework to work with high-throughput sequencing data. Bioinformatics, 31 (2), 166–169.

5. Asghar, W. & Kataoka, R. (2021) Effect of co-application of *Trichoderma* spp. with organic composts on plant growth enhancement, soil enzymes and fungal community in soil. Archives of Microbiology, 203(7), 4281–4291.

6. Atencio, L.A., Salazar, J.S., Lauter, A.N.M., Gonzales, M.D., O’Rourke, J.A. & Graham, M.A. (2021) Characterizing short- and long-term Fe stress responses in Fe deficiency tolerant and susceptible soybean (*Glycine max* L. Merr.). Plant Stress, 2, 100012.

7. Bapaume, L. & Reinhardt, D. (2012) How membranes shape plant symbioses: signaling and transport in nodulation and arbuscular mycorrhiza. Frontiers in Plant Science, 3, 223.

8. Birt, H. W. G., Tharp, C. L., Custer, G. F. & Dini-Andreote, F. (2022) Root phenotypes as modulators of microbial microhabitats. Frontiers in Plant Science, 13, 1003868.

9. Bhardwaj, D., Ansari, M.W., Sahoo, R.K. & Tuteja, N. (2014) Biofertilizers function as key players in sustainable agriculture by improving soil fertility, plant tolerance, and crop productivity. Microbial Cell Factories, 13, 66.

10. Bhat, M. A., Mishra, A. K., Shah, S. N., Bhat, M. A., Jan, S., Rahman, S., Baek, K. H. & Jan, A. T. (2024) Soil and Mineral Nutrients in Plant Health: A Prospective Study of Iron and Phosphorus in the Growth and Development of Plants. Current Issues in Molecular Biology, 46(6), 5194–5222.

11. Bjørk, P. K., Johansen, N. T., Havshøi, N. W., Rasmussen, S. A., Ipsen, J. Ø., Isbrandt, T., Larsen, T. O. & Fuglsang, A. T. (2023) *Trichoderma harzianum* peptaibols stimulate plant plasma membrane H^+^-ATPase activity. ACS Omega, 8(38), 34928–34937.

12. Blaskó, L. (2011). Alkalinity, physical effects on soils. In J. Gliński, J. Horabik, & J. Lipiec (Eds.), Encyclopedia of agrophysics (pp. 20–24). Springer. 10.1007/978-90-481-3585-1_13

13. Boeglin, L., Morère Le-Paven, M. C., Clochard, T., Fustec, J. & Limami, A. M. (2022) *Pisum sativum* response to nitrate as affected by *Rhizobium leguminosarum*-derived signals. Plants, 11(15), 1966.

14. Bolger, A.M., Lohse, M. & Usadel, B. (2014) Trimmomatic: a flexible trimmer for Illumina Sequence Data. Bioinforma. btu170.

15. Boonen, M., Mertens, J., Michiels, J. & Smolders, E. (2010) Quantitative PCR assays to enumerate *Rhizobium leguminosarum* strains in soil also target non-viable cells and overestimate those detected by the plant infection method. Soil Biology and Biochemistry, 42(12), 2342–2344

16. Borevitz, J.O.; Xia, Y.J.; Blount, J.; Dixon, R.A. & (2000) Lamb, C. Activation tagging identifies a conserved MYB regulator of phenylpropanoid biosynthesis. Plant Cell, 12, 2383–2393.

17. Brear, E. M., Day, D. A. & Smith, P. M. (2013) Fe: an essential micronutrient for the legume-rhizobium symbiosis. Frontiers in Plant Science, 4, 359.

18. Bucher, M., Hause B., Krajinski F. & Küster H. (2014). Through the doors of perception to function in arbuscular mycorrhizal symbioses. The New Phytologist, 204, 833–840.

19. Callahan, B. J., McMurdie, P. J., Rosen, M. J., Han, A. W., Johnson, A. J. & Holmes, S. P. (2016) DADA2: High-resolution sample inference from Illumina amplicon data. Nature Methods, 13(7), 581–583.

20. Canarini, A., Kaiser, C., Merchant, A., Richter, A. & Wanek, W. (2019) Root exudation of primary metabolites: mechanisms and their roles in plant responses to environment stimuli. Frontiers in Plant Science, 10, 157.

21. Chanclud, E. & Morel, J. B. (2016) Plant hormones: a fungal point of view. Molecular Plant Pathology, 17(8), 1289–1297.

22. Chaudhari, D., Rangappa, K., Das, A., Layek, J., Basavaraj, S., Kandpal, B.K., Shouche, Y., & Rahi, P. (2020) Pea (*Pisum sativum* L.) plant shapes its rhizosphere microbiome for nutrient uptake and stress amelioration in acidic soils of the North-East Region of India. Frontiers in Microbiology, 3, 968.

23. Clarke, J.D. (2009) Cetyltrimethyl ammonium bromide (CTAB) DNA miniprep for plant DNA isolation. Cold Spring Harb. Protoc., pdb.prot5177

24. Contreras-Cornejo, H. A., Schmoll, M., Esquivel-Ayala, B. A., González-Esquivel, C. E., Rocha-Ramírez, V., & Larsen, J. (2024). Mechanisms for plant growth promotion activated by Trichoderma in natural and managed terrestrial ecosystems. Microbiological Research, 281, 127621.

25. Contreras-Cornejo, H. A., Macías-Rodríguez, L., Cortés-Penagos, C., & López-Bucio, J. (2009). Trichoderma virens, a plant beneficial fungus, enhances biomass production and promotes lateral root growth through an auxin-dependent mechanism in Arabidopsis. Plant Physiology, 149(3), 1579–1592.

26. Crowley, D.E. (2006) Microbial siderophores in the plant rhizosphere. In: Barton L.L., Abadia J., editors. Fe Nutrition in Plants and Rhizospheric Microorganisms. Springer; Dordrecht, The Netherlands: pp. 169–198.

27. Curie, C. & Mari, S. (2017) New routes for plant Fe mining. The New Phytologist, 214(2), 521– 525.

28. da Costa, S.D.A., Cardoso, A.F., de Castro, G.L.S., Junior, D.D.S., da Silva, T.C. & da Silva, G.B. (2022) Co-inoculation of *Trichoderma asperellum* with *Bacillus subtilis* to promote growth and nutrient absorption in Marandu Grass. Applied and Environmental Soil Science, 10.1155/2022/3228594

29. Dhaliwal, S. S., Naresh, R. K., Mandal, A., Singh, R., & Dhaliwal, M. K. (2019). Dynamics and transformations of micronutrients in agricultural soils as influenced by organic matter build-up: A review. Environmental and Sustainability Indicators, 1–2, 100007.

30. Dasgupta, K., Ganesan, S., Manivasagam, S. & Ayre, B. G. (2011) A cytochrome P450 monooxygenase commonly used for negative selection in transgenic plants causes growth anomalies by disrupting brassinosteroid signaling. BMC Plant Biology, 11, 67.

31. Dixon, R. & Kahn, D. (2004) Genetic regulation of biological nitrogen fixation. Nature Reviews Microbiology, 2, 621–631

32. Dong, W. & Song, Y. (2020) The Significance of flavonoids in the process of biological nitrogen fixation. International Journal of Molecular Sciences, 21(16), 5926.

33. Ferrarezi, R. S., Lin, X., Gonzalez Neira, A. C., Tabay Zambon, F., Hu, H., Wang, X., Huang, J. H., & Fan, G. (2022). Substrate pH influences the nutrient absorption and rhizosphere microbiome of huanglongbing-affected grapefruit plants. Frontiers in Plant Science, 13, 856937.

34. Frawley, E. R. & Fang, F. C. (2014). The ins and outs of bacterial iron metabolism. Molecular Microbiology, 93(4), 609–616.

35. Gao, P., Qi, K., Han, Y., Ma, L., Zhang, B., Zhang, Y., Guan, X. & Qi, J. (2023) Effect of *Trichoderma viride* on rhizosphere microbial communities and biocontrol of soybean root rot. Frontiers in Microbiology, 14, 1204688.

36. García-Śanchez, M., Palma, J.M., Ocampo, J.A., García-Romera, I. & Aranda, E. (2014) Arbuscular mycorrhizal fungi alleviate oxidative stress induced by ADOR and enhance antioxidant responses of tomato plants. Journal of Plant Physiology, 171 (6), 421–428.

37. Ge, S.X., Jung, D. & Yao, R. (2020) ShinyGO: a graphical gene-set enrichment tool for animals and plants. Bioinformatics, 36 (8), 2628–2629.

38. Giovannetti, M., Binci, F., Navazio, L. & Genre, A. (2024) Nonbinary fungal signals and calcium-mediated transduction in plant immunity and symbiosis. The New Phytologist, 241(4), 1393–1400.

39. Goicoechea, N., Merino, S., & Sanchez-Diaz, M. (2004) Contribution of arbuscular mycorrhizal fungi (AMF) to the adaptations exhibited by the deciduous shrub *Anthyllis cytisoides* L. under water deficit. Plant Physiology, 122, 453–464.

40. Gorzelak, M.A., Asay, A.K., Pickles, B.J. & Simard, S.W. (2015) Inter-plant communication through mycorrhizal networks mediates complex adaptive behaviour in plant communities. AoB Plants, 7, plv050.

41. Grace, S.C. & Logan, B.A. (2000) Energy dissipation and radical scavenging by the plant phenylpropanoid pathway. Philosophical Transactions of the Royal Society B, 355, 1499– 1510.

42. Guzmán, M.G, Cellini, F., Fotopoulos, V., Balestrini, R. & Arbona, V. (2022) New approaches to improve crop tolerance to biotic and abiotic stresses. Physiologia Plantarum, 174(1), e13547.

43. Habiyaremye, J. D., Goldmann, K., Reitz, T., Herrmann, S. & Buscot, F. (2020) Tree root zone microbiome: exploring the magnitude of environmental conditions and host tree impact. Frontiers in Microbiology, 11, 749.

44. Hacisalihoglu, G., Beisel, N. S. & Settles, A. M. (2021) Characterization of pea seed nutritional value within a diverse population of *Pisum sativum*. PloS one, 16(11), e0259565.

45. Halliwell, B. (2006) Reactive species and antioxidants. Redox biology is a fundamental theme of aerobic life. Plant Physiology, 141 (2), 312–322.

46. Han, J. R., Li, S., Li, W. J., & Dong, L. (2024). Mining microbial and metabolic dark matter in extreme environments: a roadmap for harnessing the power of multi-omics data. Advanced Biotechnology, 2(3), 26.

47. Hao, Z., Xie, W., Jiang, X., Wu, Z., Zhang, X. & Chen, B. (2019) Arbuscular mycorrhizal fungus improves *Rhizobium*–Glycyrrhiza seedling symbiosis under drought stress. Agronomy, 9(10), 572.

48. Haque, A. F. M. M., Rahman, M. A., Das, U., Rahman, M. M., Elseehy, M. M., El-Shehawi, A. M., Parvez, M. S. & Kabir, A. H. (2022) Changes in physiological responses and MTP (metal tolerance protein) transcripts in soybean (*Glycine max*) exposed to differential Fe availability. Plant Physiology and Biochemistry, 179, 1–9.

49. Harbort, C. J., Hashimoto, M., Inoue, H., Niu, Y., Guan, R., Rombolà, A. D., Kopriva, S., Voges, M. J. E. E. E., Sattely, E. S., Garrido-Oter, R. & Schulze-Lefert, P. (2020) Root-secreted coumarins and the microbiota interact to improve Fe nutrition in Arabidopsis. Cell Host & Microbe, 28(6), 825–837.e6.

50. Harman, G. E., Howell, C. R., Viterbo, A., Chet, I. & Lorito, M. (2004) *Trichoderma* species-opportunistic, avirulent plant symbionts. Nature Reviews Microbiology, 2(1), 43–56.

51. Hider, R.C. & Kong, X. (2010) Chemistry and biology of siderophores. Natural Product Reports, 27 (5), 637–657.

52. Himpsl, S.D. & Mobley, H.L.T. (2019) Siderophore detection using chrome azurol S and cross-feeding assays. Methods in Molecular Biology, 2021, 97–108.

53. Hu, K. (2021) Become competent in generating RNA-seq heat maps in one day for novices without prior R experience. Methods in Molecular Biology, 2239, 269–303.

54. Huang, H., Ullah, F., Zhou, D. X., Yi, M. & Zhao, Y. (2019) Mechanisms of ROS Regulation of Plant Development and Stress Responses. Frontiers in Plant Science, 10, 800.

55. Jan, R., Kim, N., Lee, S. H., Khan, M. A., Asaf, S., Lubna, Park J. R., Asif, S., Lee, I. J. & Kim, K. M. (2021) Enhanced favonoid accumulation reduces combined salt and heat stress through regulation of transcriptional and hormonal mechanisms. Frontiers in Plant Science, 12, 796956.

56. Jin, C.W., Ye, Y.Q. & Zheng, S.J. (2014) An underground tale: contribution of microbial activity to plant iron acquisition via ecological processes. Annals of Botany, 113(1), 7–18.

57. Jin, C. W., Li, G. X., Yu, X. H., and Zheng, S. J. (2010) Plant Fe status affects the composition of siderophore-secreting microbes in the rhizosphere. Annals of Botany, 105(5), 835–841.

58. Kabir, A. H. & Bennetzen, J. L. (2024) Molecular insights into the mutualism that induces iron deficiency tolerance in sorghum inoculated with *Trichoderma harzianum*. Microbiological Research, 281, 127630.

59. Kabir, A. H., Paltridge, N. G., Able, A. J., Paull, J. G. & Stangoulis, J. C. (2012) Natural variation for Fe-efficiency is associated with upregulation of Strategy I mechanisms and enhanced citrate and ethylene synthesis in *Pisum sativum* L. Planta, 235(6), 1409–1419.

60. Kabir, A. H., Rahman, M. A., Rahman, M. M., Brailey-Jones, P., Lee, K. W. & Bennetzen, J. L. (2022) Mechanistic assessment of tolerance to Fe deficiency mediated by *Trichoderma harzianum* in soybean roots. Journal of Applied Microbiology, 133(5), 2760–2778.

61. Kabir, A.H., Rahman, M.M., Haider, S.A. & Paul, N.K. (2015) Mechanisms associated with differential tolerance to Fe deficiency in okra (*Abelmoschus escu*lentus Moench). Environmental and Experimental Botany, 112, 16–26.

62. Kim, D., Langmead, B. & Salzberg, S.L. (2015) HISAT: a fast spliced aligner with low memory requirements. Nature Methods, 12 (4), 357–360.

63. Kroh, G. E. & Pilon, M. (2019) Connecting the negatives and positives of plant Fe homeostasis. The New Phytologist, 223(3), 1052–1055.

64. Li, N., Islam, M. T. & Kang, S. (2019) Secreted metabolite-mediated interactions between rhizosphere bacteria and *Trichoderma* biocontrol agents. PloS one, 14(12), e0227228.

65. Li, S. S., Chang, Y., Li, B., Shao, S. L. & Zhen-Zhu-, Z. (2020) Functional analysis of 4-coumarate: CoA ligase from Dryopteris fragrans in transgenic tobacco enhances lignin and flavonoids. Genetics and Molecular Biology, 43(2), e20180355.

66. Liao, D., Wang, S., Cui, M., Liu, J., Chen, A. & Xu, G. (2018) Phytohormones regulate the development of arbuscular mycorrhizal symbiosis. International Journal of Molecular Sciences, 19(10), 3146.

67. Lidoy, J., Berrio, E., García, M., España-Luque, L., Pozo, M.J., & López-Ráez, J.A. (2023) Flavonoids promote *Rhizophagus irregularis* spore germination and tomato root colonization: A target for sustainable agriculture. Frontiers in Plant Science, 5, 1094194.

68. Liu, A., Zhu, Y., Wang, Y., Wang, T., Zhao, S., Feng, K., Li, L. & Wu, P. (2023) Molecular identification of phenylalanine ammonia lyase-encoding genes *EfPALs* and *EfPAL2*-interacting transcription factors in Euryale ferox. Frontiers in Plant Science, 14, 1114345.

69. Liu, H., Senthilkumar, R., Ma, G., Zou, Q., Zhu, K., Shen, X., Tian, D., Hua, M. S., Oelmüller, R. & Yeh, K. W. (2019) *Piriformospora indica*-induced phytohormone changes and root colonization strategies are highly host-specific. Plant Signaling and Behavior, 14(9), 1632688.

70. Livak, K.J. & Schmittgen, T.D. (2001) Methods. Vol. 25. San Diego, CA: Analysis of relative gene expression data using real-time quantitative PCR and the 2(-Delta Delta C(T)) Method; pp. 402–408.

71. López-Bucio, J., Cruz-Ramírez, A. & Herrera-Estrella L. (2003) The role of nutrient availability in regulating root architecture. Current Opinion in Plant Biology, 6(3), 280–287.

72. Lopez-Obando, M., Ligerot, Y., Bonhomme, S., Boyer F.D. & Rameau, C. (2015) Strigolactone biosynthesis and signaling in plant development. Development, 142, 3615–3619.

73. Lurthy, T., Cantat, C., Jeudy, C., Declerck, P., Gallardo, K., Barraud, C., Leroy, F., Ourry, A., Lemanceau, P., Salon, C. & Mazurier, S. (2020) Impact of bacterial siderophores on Fe status and ionome in Pea. Frontiers in Plant Science, 11, 730.

74. MacLean, A. M., Bravo, A. & Harrison, M. J. (2017) Plant signaling and metabolic pathways enabling arbuscular mycorrhizal symbiosis. The Plant Cell, 29(10), 2319–2335.

75. Mastouri, F., Bjorkman, T. & Harman, G.E. (2012) *Trichoderma harzianum* enhances antioxidant defense of tomato seedlings and resistance to water deficit. Molecular Plant-Microbe Interactions, 25 (9), 1264–1271.

76. McMurdie, P. J. & Holmes, S. (2013) phyloseq: an R package for reproducible interactive analysis and graphics of microbiome census data. PloS One, 8(4), e61217.

77. Meena, K. K., Sorty, A. M., Bitla, U. M., Choudhary, K., Gupta, P., Pareek, A., Singh, D. P., Prabha, R., Sahu, P. K., Gupta, V. K., Singh, H. B., Krishanani, K. K. & Minhas, P. S. (2017) Abiotic Stress Responses and Microbe-Mediated Mitigation in Plants: The Omics Strategies. Frontiers in Plant Science, 8, 172.

78. Mnguni, F. C., Padayachee, T., Chen, W., Gront, D., Yu, J. H., Nelson, D. R. & Syed, K. (2020) More P450s are involved in secondary metabolite biosynthesis in *Streptomyces* compared to *Bacillus*, Cyanobacteria, and Mycobacterium. International Journal of Molecular Sciences, 21(13), 4814.

79. Molnár, Z., Solomon, W., Mutum, L. & Janda, T. (2023) Understanding the mechanisms of Fe deficiency in the rhizosphere to promote plant resilience. Plants, 12(10), 1945.

80. Morán-Diez, M. E., Trushina, N., Lamdan, N. L., Rosenfelder, L., Mukherjee, P. K., Kenerley, C. M. & Horwitz, B. A. (2015) Host-specific transcriptomic pattern of *Trichoderma virens* during interaction with maize or tomato roots. BMC Genomics, 16(1), 8.

81. Murchie, E.H. & Niyogi, K.K. (2011) Manipulation of photoprotection to improve plant photosynthesis. Plant Physiology, 155, 86–92.

82. Nadarajah, K. K. (2020) ROS homeostasis in abiotic stress tolerance in plants. International Journal of Molecular Sciences, 21(15), 5208.

83. Nakabayashi, R., Yonekura-Sakakibara, K., Urano, K., Suzuki, M., Yamada, Y., Nishizawa, T., Matsuda, F., Kojima, M., Sakakibara, H., Shinozaki, K., Michael, A. J., Tohge, T., Yamazaki, M. & Saito, K. (2014) Enhancement of oxidative and drought tolerance in Arabidopsis by overaccumulation of antioxidant flavonoids. The Plant Journal, 77(3), 367– 379.

84. Ossler, J. N., Zielinski, C. A. & Heath, K. D. (2015) Tripartite mutualism: facilitation or trade-offs between rhizobial and mycorrhizal symbionts of legume hosts. American Journal of Botany, 102(8), 1332–1341.

85. Paffrath, V., Tandron Moya, Y. A., Weber, G., von Wirén, N. & Giehl, R. F. H. (2024) A major role of coumarin-dependent ferric Fe reduction in strategy I-type Fe acquisition in Arabidopsis. The Plant Cell, 36(3), 642–664.

86. Pandey, P.K., Bhowmik, P. & Kagale, S. (2022) Optimized methods for random and targeted mutagenesis in field pea (*Pisum sativum* L.). Frontiers in Plant Science, 8, 995542.

87. Peck, M. C., Fisher, R. F. & Long, S. R. (2006) Diverse flavonoids stimulate NodD1 binding to nod gene promoters in *Sinorhizobium meliloti*. Journal of Bacteriology, 188, 5417–5427.

88. Piyanete, C., Meechai, P. & Nakbanpotecc, W. (2009) Antioxidant activities and phenolic contents of extracts from *Salvinia olesta* and *Eichorniacrassipes*. Research Journal of Biological Sciences, 4, 1113–1117.

89. Porcel, R., Barea, J.M. & Ruiz-Lozano, J.M. (2003) Antioxidant activities in mycorrhizal soybean plants under drought stress and their possible relationship to the process of nodule senescence. The New Phytologist, 157, 135–143.

90. Poveda, J. & Eugui, D. (2022) Combined use of *Trichoderma* and beneficial bacteria (mainly *Bacillus* and *Pseudomonas*): Development of microbial synergistic bio-inoculants in sustainable agriculture. Biological Control, 176, 105100.

91. Qi, W. & Zhao, L. (2013) Study of the siderophore producing *Trichoderma asperellum* Q1 on cucumber growth promotion under salt stress. Journal of Basic Microbiology, 53(4), 355– 364.

92. Rai, S., Singh, P. K., Mankotia, S., Swain, J., & Satbhai, S. B. (2021). Iron homeostasis in plants and its crosstalk with copper, zinc, and manganese. Plant Stress, 1, 100008.

93. Ren, X., Branà, M. T., Haidukowski, M., Gallo, A., Zhang, Q., Logrieco, A. F., Li, P., Zhao, S., & Altomare, C. (2022). Potential of *Trichoderma* spp. for biocontrol of aflatoxin-producing *Aspergillus flavus*. Toxins, 14(2), 86.

94. Ren, C.G., Bai, Y.J., Kong, C.C., Bian, B. & Xie, Z.H. (2016) Synergistic interactions between salt-tolerant rhizobia and arbuscular mycorrhizal fungi on salinity tolerance of *Sesbania cannabina* plants. Journal of Plant Growth Regulation, 35, 1098–1107.

95. Rodríguez-Celma, J. and Schmidt, W. (2013) Reduction-based Fe uptake revisited: on the role of secreted Fe-binding compounds. Plant Signal Behav, 8, e26116.

96. Russo, G., Carotenuto, G., Fiorilli, V., Volpe, V., Chiapello, M., Van Damme, D. & Genre, A. (2019) Ectopic activation of cortical cell division during the accommodation of arbuscular mycorrhizal fungi. The New Phytologist, 221(2), 1036–1048.

97. Saleem, A., Zulfiqar, A., Saleem, M. Z., Ali, B., Saleem, M. H., Ali, S., Tufekci, E. D., Tufekci, A. R., Rahimi, M., & Mostafa, R. M. (2023). Alkaline and acidic soil constraints on iron accumulation by Rice cultivars in relation to several physio-biochemical parameters. BMC Plant Biology, 23(1), 397.

98. Santiago, A., Quintero, J.M., Aviles, M. & Delgado, A. (2011) Effect of *Trichoderma asperellum* strain T34 on Fe, copper, manganese, and zinc uptake by wheat grown on a calcareous medium. Plant and Soil, 342, 97–104.

99. Schenkeveld, W. D., Reichwein, A. M., Bugter, M. H., Temminghoff, E. J., & van Riemsdijk, W. H. (2010) Performance of soil-applied FeEDDHA isomers in delivering Fe to soybean plants in relation to the moment of application. Journal of Agricultural and Food Chemistry, 58(24), 12833–12839.

100. Schliemann, W., Ammer, C. & Strack, D. (2008) Metabolite profiling of mycorrhizal roots of *Medicago truncatula*. Phytochemistry, 69(1), 112–146.

101. Shumilina, J., Soboleva, A., Abakumov, E., Shtark, O. Y., Zhukov, V. A. & Frolov, A. (2023) Signaling in Legume-Rhizobia Symbiosis. International Journal of Molecular Sciences, 24(24), 17397.

102. Singh, D., Geat, N., Jadon, K. S., Verma, A., Sharma, R., Rajput, L. S., Mahla, H. R., & Kakani, R. K. (2024). Isolation and characterization of biocontrol microbes for development of effective microbial consortia for managing *Rhizoctonia bataticola* root rot of cluster bean under hot arid climatic conditions. Microorganisms, 12(11), 2331.

103. Singh, S.P., Pandey, S., Mishra, N., Giri, V.P., Mahfooz, S., Bhattacharya, A., Kumari, M., Chauhan, P., Verma, P., Nautiyal, C.S. & Mishra, A. (2019) Supplementation of *Trichoderma* improves the alteration of nutrient allocation and transporter genes expression in rice under nutrient deficiencies. Plant Physiology and Biochemistry, 143, 351–363.

104. Staropoli, A., Di Mola, I., Ottaiano, L., Cozzolino, E., Pironti, A., Lombardi, N., Nanni, B., Mori, M., Vinale, F., Woo, S. L., & Marra, R. (2024). Biodegradable mulch films and bioformulations based on *Trichoderma* sp. and seaweed extract differentially affect the metabolome of industrial tomato plants. Journal of fungi, 10(2), 97.

105. Stringlis, I. A., de Jonge, R. & Pieterse, C. M. J. (2019) The age of coumarins in plant-microbe Interactions. Plant and Cell Physiology, 60(7), 1405–1419.

106. Sun, C., Wu, T., Zhai, L., Li, D., Zhang, X., Xu, X. & Han, Z. (2016) Reactive oxygen species function to mediate the Fe deficiency response in an Fe-efficient apple Cultivar: An early response mechanism for enhancing reactive oxygen production. Frontiers in Plant Science, 7, 1726.

107. Tewari, R.K., Hadacek, F., Sassmann, S. & Lang, I. (2013) Fe deprivation-induced reactive oxygen species generation leads to non-autolytic PCD in *Brassica napus* leaves. Environmental and Experimental Botany, 91, 74–83

108. Thilakarathna, M. S. & Cope, K. R. (2021) Split-root assays for studying legume-rhizobia symbioses, rhizodeposition, and belowground nitrogen transfer in legumes. Journal of Experimental Botany, 72(15), 5285–5299.

109. Timofeeva, A. M., Galyamova, M. R. & Sedykh, S. E. (2022) Bacterial siderophores: classification, biosynthesis, perspectives of use in agriculture. Plants, 11(22), 3065.

110. Tsai, H. H., Wang, J., Geldner, N. & Zhou, F. (2023) Spatiotemporal control of root immune responses during microbial colonization. Current Opinion in Plant Biology, 74, 102369.

111. Uddling, J., Gelang-Alfredsson, J., Piikki, K. & Pleijel, H. (2007) Evaluating the relationship between leaf chlorophyll concentration and SPAD-502 chlorophyll meter readings. Photosynthesis Research, 91(1), 37–46.

112. Umadevi, P., Anandaraj, M., Srivastav, V. & Benjamin, S. (2018) *Trichoderma harzianum* MTCC 5179 impacts the population and functional dynamics of microbial community in the rhizosphere of black pepper (*Piper nigrum* L.). Brazilian Journal of Microbiology, 49(3), 463–470.

113. Vert, G., Briat, J.F. & Curie, C. (2001) Arabidopsis *IRT2* gene encodes a root-periphery Fe transporter. The Plant Journal, 26, 181–189.

114. Vinale, F., Nigro, M., Sivasithamparam, K., Flematti, G., Ghisalberti, E.L., Ruocco, M. & Lorito, M. (2013) Harzianic acid: a novel siderophore from *Trichoderma harzianum*. FEMS Microbiology Letters, 347 (2), 123–129.

115. Vukelić, I. D., Prokić, L. T., Racić, G. M., Pešić, M. B., Bojović, M. M., Sierka, E. M., Kalaji, H. M., & Panković, D. M. (2021). Effects of *Trichoderma harzianum* on photosynthetic characteristics and fruit quality of tomato plants. International Journal of Molecular Sciences, 22(13), 6961.

116. Wang, N., Wang, T., Chen, Y., Wang, M., Lu, Q., Wang, K., Dou, Z., Chi, Z., Qiu, W., Dai, J., Niu, L., Cui, J., Wei, Z., Zhang, F., Kümmerli, R. & Zuo, Y. (2024) Microbiome convergence enables siderophore-secreting-rhizobacteria to improve Fe nutrition and yield of peanut intercropped with maize. Nature Communications, 15(1), 839.

117. Wang, L., Chen, M., Lam, P. Y., Dini-Andreote, F., Dai, L., & Wei, Z. (2022). Multifaceted roles of flavonoids mediating plant-microbe interactions. Microbiome, 10(1), 233.

118. Wang, Q., Liu, J. & Zhu, H. (2018) Genetic and molecular mechanisms underlying symbiotic specificity in legume-Rhizobium interactions. Frontiers in Plant Science, 9, 313.

119. Wiesmann, C.L., Wang, N.R., Zhang, Y., Liu, Z. & Haney, C.H. (2023) Origins of symbiosis: shared mechanisms underlying microbial pathogenesis, commensalism and mutualism of plants and animals. FEMS Microbiology Reviews, 47, fuac048.

120. Wu, H., Zou, Q., Luo, S., He, D., Li, X., Yan, C. & Cheng, G. (2020) Lack of the slavin-containing monooxygenase FmoA partially impairs the symbiotic interaction of *Mesorhizobium Huakuii* with *Astragalus Sinicus*. BMC Microbiology, 10.21203/rs.3.rs-27635/v1

121. Xie, W., Hao, Z.P., Zhou, X.F., Jiang, X.L., Xu, L.J., Wu, S.L., Zhao, A.H., Zhang, X. & Chen, B.D. (2018) Arbuscular mycorrhiza facilitates the accumulation of glycyrrhizin and liquiritin in *Glycyrrhiza uralensis* under drought stress. Mycorrhiza, 28, 285–300.

122. Xiong, H., Hua, L., Reyna-Llorens, I., Shi, Y., Chen, K.M., Smirnoff, N. & Hibberd, J.M. (2021) Photosynthesis-independent production of reactive oxygen species in the rice bundle sheath during high light is mediated by NADPH oxidase. Proceedings of the National Academy of Sciences of the United States of America, 118, e2022702118

123. Yadav, R. K., Tripathi, M. K., Tiwari, S., Tripathi, N., Asati, R., Chauhan, S., Tiwari, P. N. & Payasi, D. K. (2023) Genome editing and improvement of abiotic stress tolerance in crop plants. Life, 13(7), 1456.

124. Yates, A. D., Allen, J., Amode, R. M., Azov, A. G., Barba, M., Becerra, A., Bhai, J., Campbell, L. I., Carbajo Martinez, M., Chakiachvili, M., Chougule, K., Christensen, M., Contreras-Moreira, B., Cuzick, A., Da Rin Fioretto, L., Davis, P., De Silva, N. H., Diamantakis, S., Dyer, S., Elser, J. & Flicek, P. (2022) Ensembl Genomes 2022: an expanding genome resource for non-vertebrates. Nucleic Acids Research, 50(D1), D996–D1003.

125. Ye, Q., Park, J.E., Gugnani, K., Betharia, S., Pino-Figueroa, A. & Kim, J. (2017) Influence of Fe metabolism on manganese transport and toxicity. Metallomics, 16, 1028–1046.

126. Zamboni, A., Zanin, L., Tomasi, N., Pezzotti, M., Pinton, R., Varanini, Z. & Cesco, S. (2012) Genome-wide microarray analysis of tomato roots showed defined responses to iron deficiency. BMC Genomics, 13, 101.

127. Zhao, L., Wang, Y. & Kong, S. (2020) Effects of *Trichoderma asperellum* and its siderophores on endogenous auxin in *Arabidopsis thaliana* under Fe-deficiency stress. International Microbiology, 23(4), 501–509.

