## Supplementary for "*Trichoderma afroharzianum* T22 induces rhizobia and flavonoid-driven symbiosis to promote tolerance to alkaline stress in garden pea"

**Supplementary Table S1.** Physico-chemical properties of soil used for the cultivation of pea plants.

| **Texture**  Clay | | | | | **Organic matter**  5% | | | | | **Cation Exchange Capacity**  15.6 meq/100g | | | | **Water Holding Capacity**  42% | | | |
| --- | --- | --- | --- | --- | --- | --- | --- | --- | --- | --- | --- | --- | --- | --- | --- | --- | --- |
| **Elemental content ppm** | | | | | | | | | | | | | | | | | |
| **Al** aluminum | **N** arsenic | **B** boron | **Ca** calcium | **Cd** cadmium | **Cr** chromium | **Cu** copper | **Fe** iron | **K** potassium | **Mg** magnesium | **Mn** manganese | **Mo** molybdenum | **Na** sodium | **Ni** nickel | **P** phosphorus | **Pb** lead | **S** sulfur | **Zn** zinc |
| no limit | no limit | no limit | no limit | <2 | <100 | <100 | no limit | no limit | no limit | <3500 | <440 | no limit | <50 | no limit | <75 | no limit | <100 |
| 7686 | 112 | 8.97 | 1415 | 0.24 | 10.2 | 11.0 | 9033 | 1286 | 1831 | 641 | 0.42 | 991 | 9.33 | 626 | 21.0 | 182 | 68.8 |

**Supplementary Table S2.** Gene-specific primers used in qPCR experiments.

| Gene | Primers |
| --- | --- |
| *PsGAPDH* | Fw GTGGTCTCCACTGACTTTATTGGT  Rv TTCCTGCCTTGGCATCAAA |
| *NifA* | Fw-TGCCGGATGTCTATCCAAAT  Rv-AGTCGCAGCGGACAGTAGAT |
| *NifD* | Fw-GGGTCGGATGCATCAAGCAAG  Rv-GTTGCTAGGCTCATACGAATATC |
| *NifH* | Fw-CGACCACGTCACAGAACAC  Rv-CCTTGTAGCCGACCTTCATG |
| *PsCoA* | Fw-TGTCTCATCGTCGCTTCCAC  Rv- AGAATGGATTGGCGGTGGTT |
| *RHIZ* | Fw- AGAGTTTGATCCTGGCTCAG  Rv- TTGACTACGGAATAACGCAG |

**Supplementary Table S3.** The top differentially expressed genes in roots of garden pea and their expression in different growth conditions. Values marked with an asterisk are significant > Log2 fold, < P value 0.05.

| Genes | Function | Alkaline+T22 vs Alkaline | Fe- vs Control | T22+ vs Alkaline+T22 |
| --- | --- | --- | --- | --- |
| Psat5g201640 | Monooxygenase activity | 2.14^*^ | -3.30^*^ | 1.17 |
| Psat3g072160 | Ammonia−lyase activity | 2.39^*^ | -2.59^*^ | 0.91 |
| Psat3g201240 | UDP−glycosyltransferase activity | 2.17^*^ | -3.26^*^ | 0.95 |
| Psat2g001960 | Ammonia−lyase activity | 3.52^*^ | -2.52 | 0.59 |
| Psat1g068400 | UDP−glycosyltransferase activity | 2.90^*^ | -4.34^*^ | 0.94 |
| Psat7g063080 | UDP−glycosyltransferase activity | 3.51^*^ | -4.21^*^ | 0.82 |
| Psat1g006320 | Auxin responsive protein | 5.70^*^ | 0.72 | 0.48 |
| Psat2g188920 | 4−coumarate:coenzyme A ligase | 2.34^*^ | -0.83 | 0.58 |
| Psat7g116000 | Monooxygenase activity | 2.79^*^ | 1.02 | 0.68 |
| Psat5g190680 | ABC−type transporter activity | 2.46^*^ | -0.14 | 0.71 |
| Psat6g141240 | Manganese ion binding | 2.49^*^ | 0.75 | 0.42 |
| Psat3g159880 | Iron ion binding | 2.10^*^ | -0.57 | 0.12 |
| Psat4g113000 | Nitrate transporter activity | 2.66^*^ | -0.44 | 0.47 |
| Psat5g286200 | Iron ion binding | 3.00^*^ | 2.86^*^ | 1.07 |
| Psat4g000120 | Inorganic phosphate transporter | 2.80^*^ | -1.92 | 0.78 |
| Psat2g189680 | Zinc ion binding | 3.02^*^ | -1.88 | 0.83 |
| Psat1g123040 | Magnesium ion binding | 2.67^*^ | -1.68 | 0.75 |
| Psat1g086520 | Oxidoreductase activity | 3.74^*^ | -2.46^*^ | 0.69 |
| Psat1g011640 | Metal ion transporter activity | 4.22^*^ | -2.31^*^ | 0.78 |
| Psat1g171760 | Metal ion binding | 3.19^*^ | -2.28 | 0.72 |
| Psat1g168800 | Peroxidase activity | 5.87^*^ | -1.15 | 0.40 |
| Psat3g121200 | Peroxidase activity | 2.10^*^ | -0.25 | 0.33 |
| Psat2g065320 | Glutamyl−tRNA reductase activity | 3.21^*^ | -2.06 | 0.76 |
| Psat3g020440 | Glutathione peroxidase activity | 6.36^*^ | -3.72^*^ | 0.55 |
| Psat5g186720 | Meristem development | 3.58^*^ | -3.38^*^ | 1.05 |
| Psat5g198360 | Glutathione S-transferase | 3.07^*^ | -1.14^*^ | 0.52 |
| Psat7g008120 | Calcium ion binding | 2.47^*^ | -4.03^*^ | 0.96 |
| Psat2g172320 | Growth factor activity | 3.71^*^ | -1.30 | 0.76 |
| Psat7g008720 | Pyrophosphatase activity | 2.42^*^ | -3.44^*^ | 0.70 |
| Psat2g173800 | Peroxidase activity | 3.19^*^ | -2.98 | 0.57 |

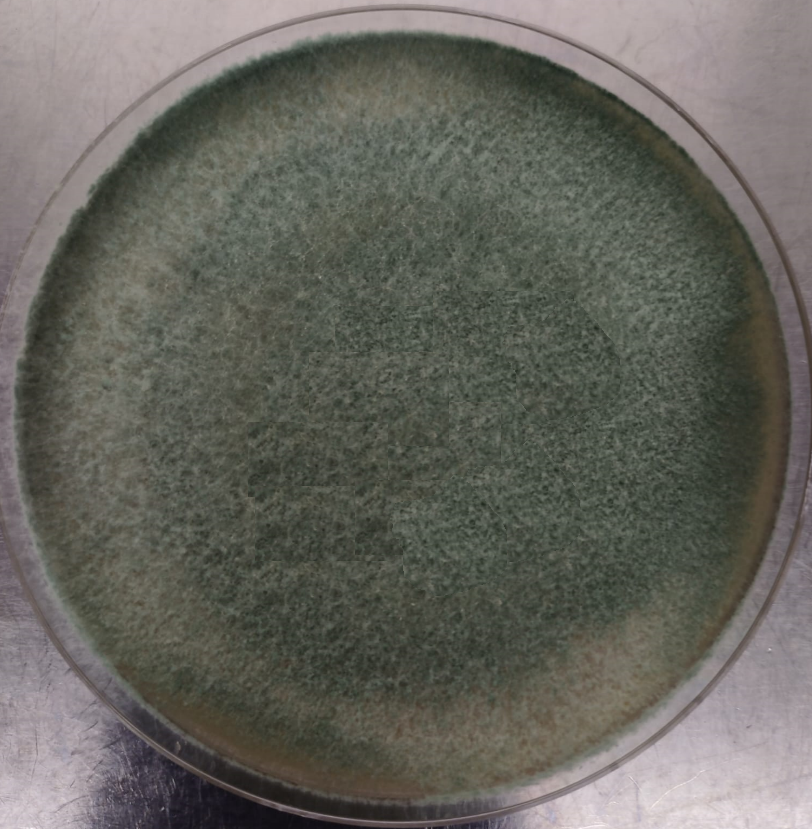

**Supplementary Fig. S1.** Culture of *T. afroharzianum* T22 inoculum in potato dextrose agar (PDA) after 3 days of incubation at 25°C.

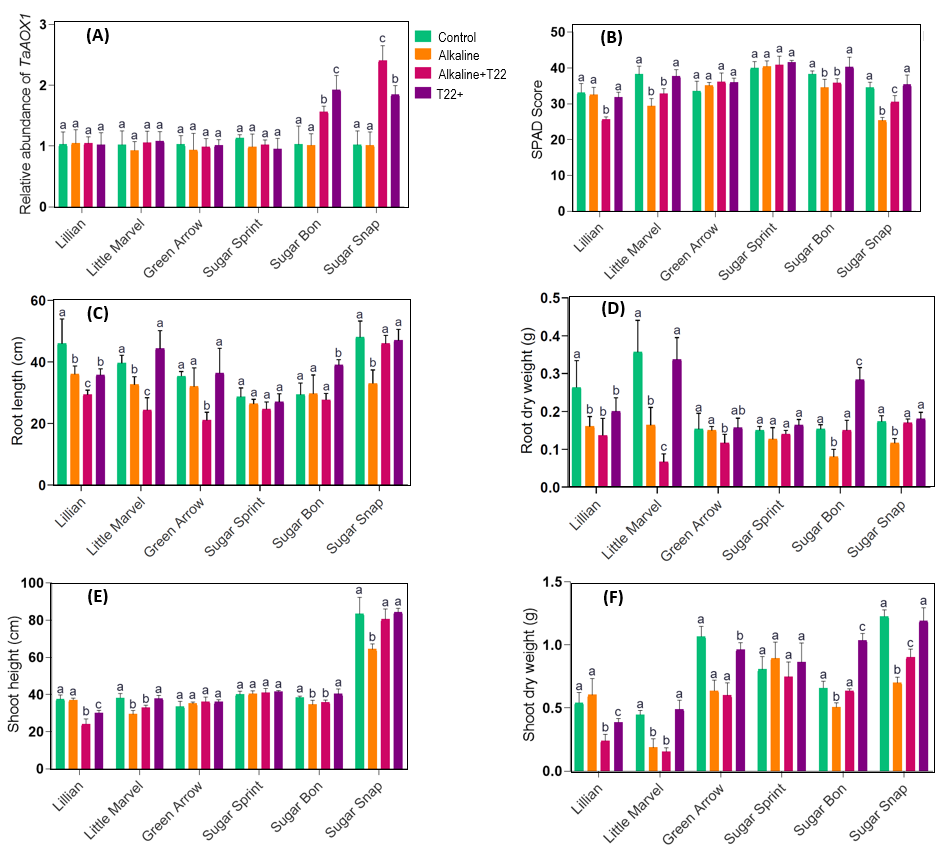

**Supplementary Fig. S2.** Screening of genetic lines that are sensitive to soil alkalinity but capable of leveraging *T. afroharzianum* T22 to combat alkaline stress: relative abundance of *TaAOX1* (A), leaf SPAD score (B), root length (C), root dry weight (D), shoot height (E) and shoot dry weight (F). The data presented are means with standard deviations (*n* = 3 individual replicates). Statistically significant differences in the ANOVA among different treatments at a *p* < 0.05 level were indicated by different alphabetical letters above the bars.

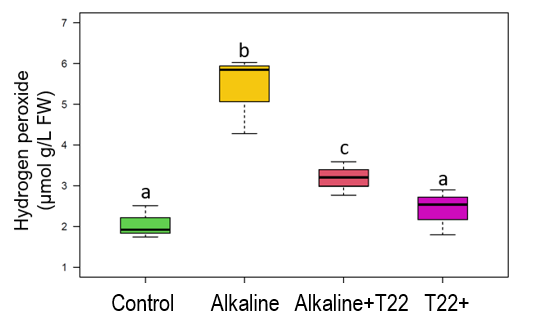

**Supplementary Fig. S3.** Concentration of hydrogen peroxide in roots of Sugar Snap cultivated in the absence or presence of alkaline stress inoculated with (T22+) or without T22. The data presented are means with standard deviations (*n* = 3 individual replicates). Statistically significant differences in the ANOVA test among different treatments (*p* < 0.05) are indicated by different letters above the bars.

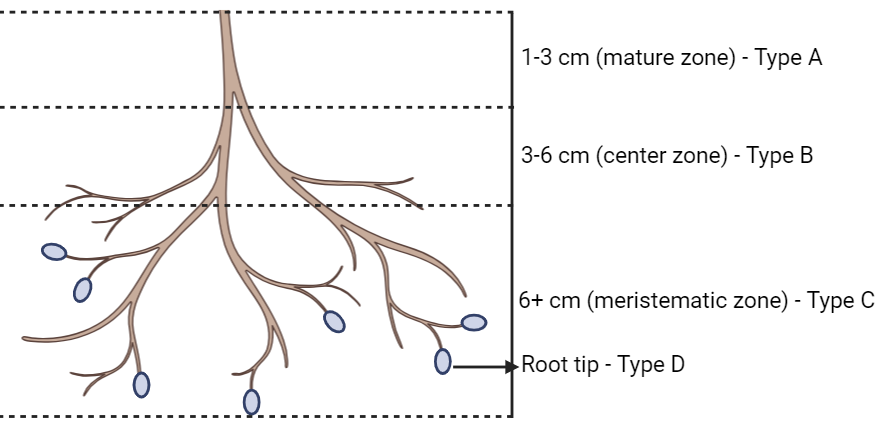

**Supplementary Fig. S4.** Schematic representation of different root partitions used for the colonization efficiency of T22.

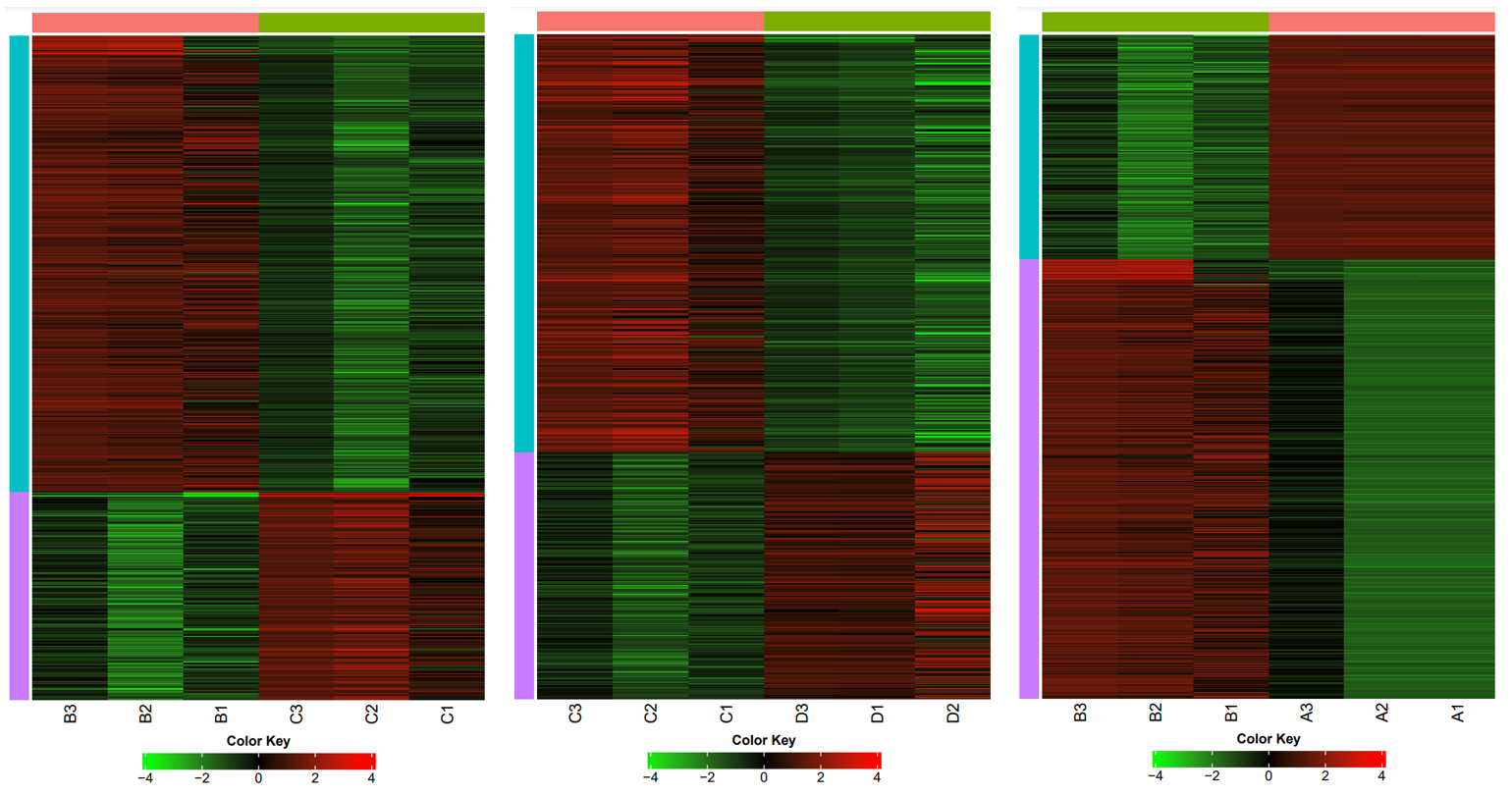

**Supplementary Fig. S5.** Heatmap of gene expression profiles showing variability based on fold change among biological replicates of different treatment groups (A: control, B: Fe-, C: Fe-T22+, and D: T22).

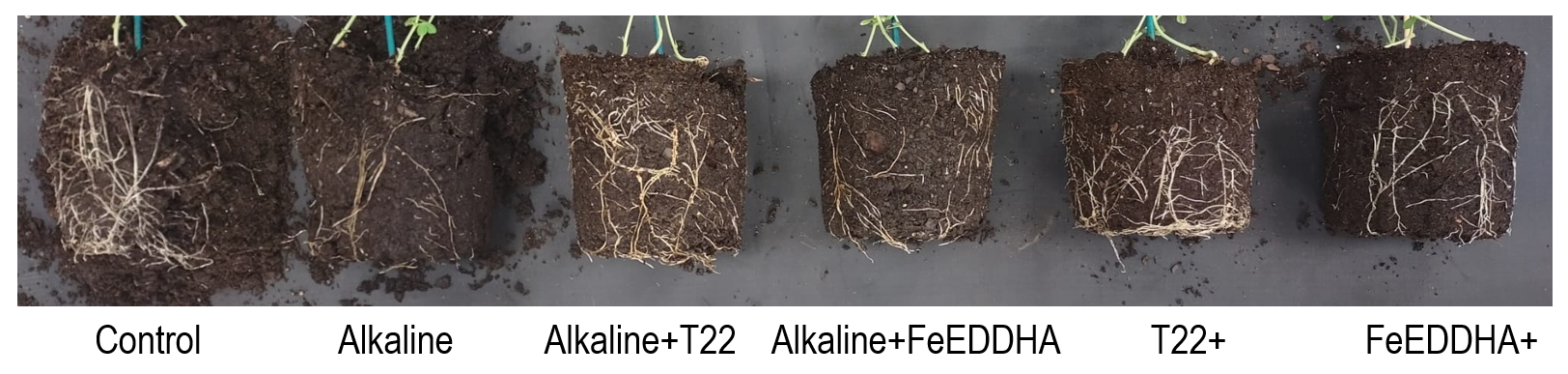

**Supplementary Fig. S6.** Effect of FeEDDHA on root development in pea plants cultivated with or without the absence (control) or presence of alkaline stress and T22.

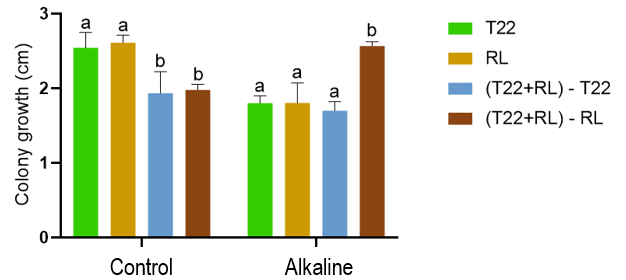

**Supplementary Fig. S7.** Co-culture of *T. afroharzianum* (T22) and *R. leguminosarum* (RL) in nutrient agar under control and alkaline conditions for 7 days. The data presented are means with standard deviations (*n* = 3 individual replicates). Statistically significant differences in the ANOVA test among different treatments (*p* < 0.05) are indicated by different letters above the bars.
